# Cytochrome c_M_ downscales photosynthesis under photomixotrophy in *Synechocystis* sp. PCC 6803

**DOI:** 10.1101/853416

**Authors:** Daniel Solymosi, Dorota Muth-Pawlak, Lauri Nikkanen, Duncan Fitzpatrick, Ravendran Vasudevan, Christopher J. Howe, David J. Lea-Smith, Yagut Allahverdiyeva

## Abstract

Photomixotrophy is a metabolic state, which enables photosynthetic microorganisms to simultaneously perform photosynthesis and metabolism of imported organic carbon substrates. This process is complicated in cyanobacteria, since many, including *Synechocystis* sp. PCC 6803, conduct photosynthesis and respiration in an interlinked thylakoid membrane electron transport chain. Under photomixotrophy, the cell must therefore tightly regulate electron fluxes from photosynthetic and respiratory complexes. In this study, we show via characterization of photosynthetic apparatus and the proteome, that photomixotrophic growth results in a gradual reduction of the plastoquinone pool in wild-type *Synechocystis*, which fully downscales photosynthesis over three days of growth. This process is circumvented by deleting the gene encoding cytochrome *c*_M_ (CytM), a cryptic *c*-type heme protein widespread in cyanobacteria. ΔCytM maintained active photosynthesis over the three day period, demonstrated by high photosynthetic O_2_ and CO_2_ fluxes and effective yields of Photosystem II and Photosystem I. Overall, this resulted in a higher growth rate than wild-type, which was maintained by accumulation of proteins involved in phosphate and metal uptake, and cofactor biosynthetic enzymes. While the exact role of CytM has not been determined, a mutant deficient in the thylakoid-localised respiratory terminal oxidases and CytM (ΔCox/Cyd/CytM) displayed a similar phenotype under photomixotrophy to ΔCytM, demonstrating that CytM is not transferring electrons to these complexes, which has previously been suggested. In summary, the obtained data suggests that CytM may have a regulatory role in photomixotrophy by reducing the photosynthetic capacity of cells.

**One sentence summary:** The cryptic, highly conserved cytochrome c_M_ completely blocks photosynthesis in *Synechocystis* under three days of photomixotrophy, possibly by suppressing CO_2_ assimilation.

## Introduction

Switching between different trophic modes is an advantageous feature, which provides great metabolic flexibility for cyanobacteria. For a long time, these photosynthetic prokaryotes were considered as a group of predominantly photoautotrophic organisms (Smith 1983, Stal and Moezelaar 1997). Lately, accumulating evidence marks the physiological and ecological importance of trophic modes involving organic carbon assimilation, *e*.*g*. photomixotrophy (Zubkov and Tarran 2008, Moore et al 2013). Dissolved organic carbon, most notably monosaccharides, glucose and fructose, accumulates in the environment, mainly during phytoplankton blooms (Teeling et al 2012, Ittekot et al 1981). During photomixotrophy, photosynthetic organisms must balance the consumption of organic carbon sources with photosynthesis and carbon fixation.

In the model cyanobacterium, *Synechocystis* sp. PCC 6803 (hereafter referred to as *Synechocystis*), photomixotrophy is further complicated by the operation of anabolic and catabolic processes occurring in the same cellular compartment and by the presence of an interlinked thylakoid membrane-localised electron transport pathway involved in both photosynthesis and respiration (Vermaas et al. 2001, Mullineaux 2014, Lea-Smith et al 2016). In *Synechocystis*, photosynthetic linear electron flow is similar to other oxygenic photoautotrophs. In photosystem (PS) II and PSI, the energy of the harvested photons induces charge separation. Electron from the PSII primary donor P680 passes via pheophytin and the primary quinone Q_A_, to the secondary quinone, Q_B_. Oxidized P680^+^ is the strongest biological oxidizing molecule, which drives water splitting on the lumenal side of PSII. When Q_B_ is doubly reduced, it binds two protons from the cytosol, converting plastoquinone (PQ) to plastoquinol (PQH_2_), then diffuses into the membrane PQ pool. Cytochrome (Cyt) *b*_6_*f* receives two electrons and transfers an electron to the mobile small protein, plastocyanin (Pc) or cytochrome c_6_ (Cyt c_6_). An electron is subsequently transferred to PSI, replacing a newly excited electron that is transferred from the PSI reaction center P700^+^ via several co-factors to ferredoxin (Fed). Lastly, electrons are transferred from Fed to NADP^+^ by ferredoxin-NADP^+^ reductase (FNR) to generate NADPH. In the respiratory electron transfer pathway, PQ is reduced by NAD(P)H dehydrogenase-like complex I (NDH-1) and succinate dehydrogenase (SDH), using electrons ultimately derived from Fed (Shuller et al 2019) and succinate, respectively. Electrons from PQ-pool can be transferred to a thylakoid-localized respiratory terminal oxidase (RTO), cytochrome *bd*-quinol oxidase (Cyd), or via Cyt *b*_6_*f* then Pc/Cyt *c*_*6*_, to a second RTO, an *aa*_*3*_-type cytochrome-*c* oxidase complex (Cox). How *Synechocystis* regulates electron input from PSII and the NDH-1 and SDH complexes into the photosynthetic electron transport chain and to respiratory terminal acceptors under photomixotrophic conditions, is not fully understood. Moreover, *Synechocystis* encodes four isoforms of flavodiiron proteins (FDPs), Flv1-4, which likely utilize NAD(P)H (Vicente et al. 2002, Brown et al 2019) or reduced Fed (Santana-Sanchez et al. 2019). These proteins function in light-induced O_2_ reduction as hetero-oligomers consisting of either Flv1/Flv3 or Flv2/Flv4 (Helman et al. 2003, Mustila et al. 2016, Allahverdiyeva et al. 2015, Santana-Sanchez et al. 2019).

In *Synechocystis*, a *c*-type Cyt, the water-soluble Cyt *c*_6_ (formerly referred to as Cyt c_553_), can substitute for Pc under conditions of copper deprivation (Durán et al 2004). Cyt *c*_6_ belongs to the Cyt *c* family, whose members are characterized by a covalently bound *c-*type heme cofactor. *C-*type Cyts are further classified into groups such as the Cyt *c*_6_-like proteins, Cyt *c*_*555*_, Cyt *c*_*550*_, and CytM (Bialek et al. 2008). Apart from the well-established role of Cyt *c*_6_ in electron transfer (Kerfeld and Krogman 1998) and the role of Cyt *c*_*550*_ (PsbV) in stabilizing the PS II water splitting complex (Shen and Inoue 1993), most of the Cyt *c* proteins remain enigmatic.

CytM is conserved in nearly every sequenced cyanobacteria with the exception of the obligate symbionts, *Candidatus acetocyanobacterium thalassa* and *Candidatus Synechococcus spongiarum* (Supplemental Fig. S1, Bialek et al 2016). In *Synechocystis*, CytM is encoded by *sll1245* (Malakhov et al 1994). Nevertheless, its subcellular location is ambiguous. An early study localised CytM to the thylakoid and plasma membranes in ‘purified’ membrane fractions (Bernroitner et al 2009). However, cross contamination between membranes was not determined, which has been an issue in studies using similar separation techniques (Sonoda et al 1997, Schultze et al 2009). In later proteomics studies, CytM has not been detected or localised using membranes purified by either two-phase aqueous polymer partitioning or subcellular fractionation (Baers et al 2019). However, the structure of the hydrophobic N-terminus resembles a signal peptide, which suggests that CytM is targeted to a membrane. Sequence similarity to the N-terminus cleavage site of *Synechocystis* Cyt *c*_6_ suggests that the N-terminus is processed and the mature 8.3 kDa protein is inserted into the lumen (Malakhov et al 1994). However, cleavage does not seem to occur *in vivo,* as the protein extracted from various cyanobacterial species, including *Synechocystis, Synechococcus elongatus* PCC 6301 and *Anabaena* sp. PCC 7120, was found to be around 12 kDa (Cho et al 2000, Bernroitner et al 2009), implying that the hydrophobic N-terminus remains on the protein and serves as a membrane anchor.

It has been suggested that CytM may play a role in respiratory or photosynthetic electron transfer (Vermaas et al 2001, Bernroitner et al 2009). In *Synechocystis*, CytM was shown to reduce the Cu_A_ center of Cox *in vitro* with similar efficiency as Cyt *c*_6_ (Bernroitner et al 2009). However, given the midpoint potential of CytM (+150 mV), electron transfer from Cyt *b*_6_*f* (+320 mV) to CytM would be energetically uphill (Cho et al 2000). Notably, CytM is unable to reduce PSI *in vitro* (Molina-Heredia et al 2002). Thus, it is difficult to see how the protein would substitute for Cyt *c*_6_ or Pc. Importantly, CytM is not detected under photoautotrophic conditions (Baers et al 2019) and deletion of the gene does not affect net photosynthesis or dark respiratory rates (Malakhov et al 1994) under these conditions. Cold, high light and salt stress, however, induce gene expression and the stress-induced co-transcriptional regulation between *cytM* (CytM), *petJ* (Cyt *c*_6_) and *petE* (Pc) suggests a stress-related role in electron transfer (Malakhov et al 1999).

Besides environmental stresses, CytM has been linked to organic carbon-assimilating trophic modes. A strain of *Leptolyngbya boryana* was found to grow faster than wild-type (WT) in dark heterotrophy. Genome re-sequencing revealed that the fast-growing strain harbours a disrupted *cytM* (Hiraide et al 2015). In line with this, the *cytM* deletion mutant of *Synechocystis* demonstrated a growth advantage over the WT under dark and light-activated heterotrophic conditions, and under photomixotrophic conditions (Hiraide et al 2015). Under dark heterotrophic conditions, ΔCytM had higher respiratory and photosynthetic activity. However, the physiological mechanism and the functional role of CytM remains entirely unknown.

In this study, we sought to uncover the physiological background behind the growth advantage of photomixotrophically grown *Synechocystis* ΔCytM by characterizing its photosynthetic machinery and the proteomic landscape. We demonstrate that a mutant lacking CytM circumvents over-reduction of the PQ-pool during photomixotrophic growth, enabling higher rates of net photosynthesis. In order to meet the substrate demand for enhanced growth, the mutant accumulates transporter proteins, cofactor biosynthetic enzymes and slightly adjusts central carbon metabolism. Although the function of CytM was previously associated with Cox, both thylakoid respiratory terminal oxidases, Cox and Cyd, were found to be dispensable for the metabolic advantage conferred by deletion of CytM in photomixotrophy. We conclude that when cells are exposed to high glucose conditions, CytM reduces the photosynthetic capacity and contributes to regulating the redox state of the intertwined photosynthetic and respiratory electron transport chain, in order to accommodate this new energy source.

## Results

### Deletion of CytM confers growth advantage on ΔCytM and ΔCox/Cyd/CytM in photomixotrophy

In order to elucidate the physiological role of CytM and its possible functional association with thylakoid-localised RTOs, we studied the ΔCytM, ΔCox/Cyd and ΔCox/Cyd/CytM mutants. Unmarked mutants of *Synechocystis* lacking CytM were constructed by disrupting the *cytM* gene (*sll1245*) in WT (Supplemental Fig. S2) and the ΔCox/Cyd mutant (Lea-smith et al. 2013). Strains were then pre-cultured under photoautotrophic conditions at 3 % CO_2_ and examined under a range of different growth conditions at air level CO_2_.

First, we determined whether deletion of *cytM* affected photoautotrophic growth by culturing cells under moderate constant 50 µmol photons m^−2^ s^−1^ light. In line with previous studies (Malakhov et al 1994, Hiraide et al 2015), no growth difference was observed between ΔCytM and WT under photoautotrophic conditions (Fig 1A).

**Figure 1.**
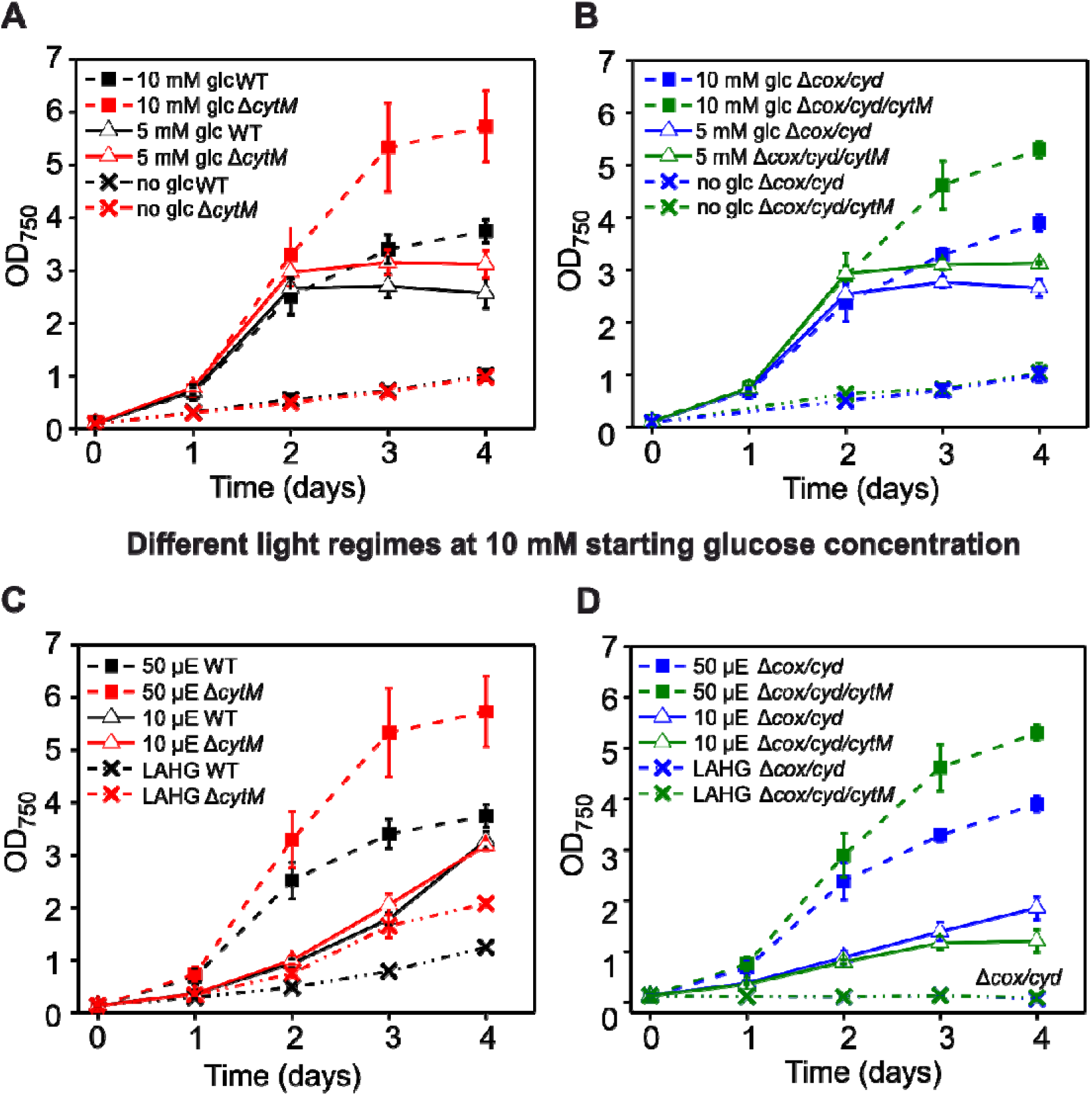
Impact of different glucose concentrations and light regimes on the growth of WT, ΔCytM, ΔCox/Cyd and ΔCox/Cyd/CytM. Cultures were exposed to 50 µmol photons m^−2^ s^−1^ light (A, B) and were grown under photoautotrophic conditions without glucose (dash-dot-dot line) or under photomixotrophic conditions with 5 mM glucose (solid line) or 10 mM glucose (dashed line). Growth was then assessed under various light regimes in cultures containing 10 mM glucose (C, D), under constant 50 µmol photons m^−2^ s^−1^ light (dashed line), constant 10 µmol photons m^−2^ s^−1^ light (solid line) and LAHG (light-activated heterotrophic growth) which included 15 min of 50 µmol photons m^−2^ s^−1^ light exposure every 24 h (dash-dot-dot line). Values are means ± SD, n = 3-7 biological repeats.

Next, we characterized growth under photomixotrophic conditions. To determine how different starting glucose concentrations affected photomixotrophic growth (Fig. 1A, B), we supplemented the medium with 5 mM and 10 mM glucose and cultivated the strains under constant 50 µmol photons m^−2^ s^−1^ light. Based on optical density measurements (OD_750_), all cultures with added glucose grew substantially faster than those cultured photoautotrophically (Fig 1A, B). Deletion of *cytM* had no effect on cells grown at 5 mM glucose. However, when cultured with 10 mM glucose, ΔCytM demonstrated 1.9±0.4 (p=6E-6) higher OD_750_ than WT and ΔCox/Cyd/CytM demonstrated 1.9±0.6 (p=0.002) higher OD_750_ compared to ΔCox/Cyd, after three days. In line with this, ΔCytM consumed more glucose than WT (Fig. 2A), as quantified by measuring the glucose concentration of the cell-free spent media on the third day of photomixotrophic growth.

**Figure 2.**
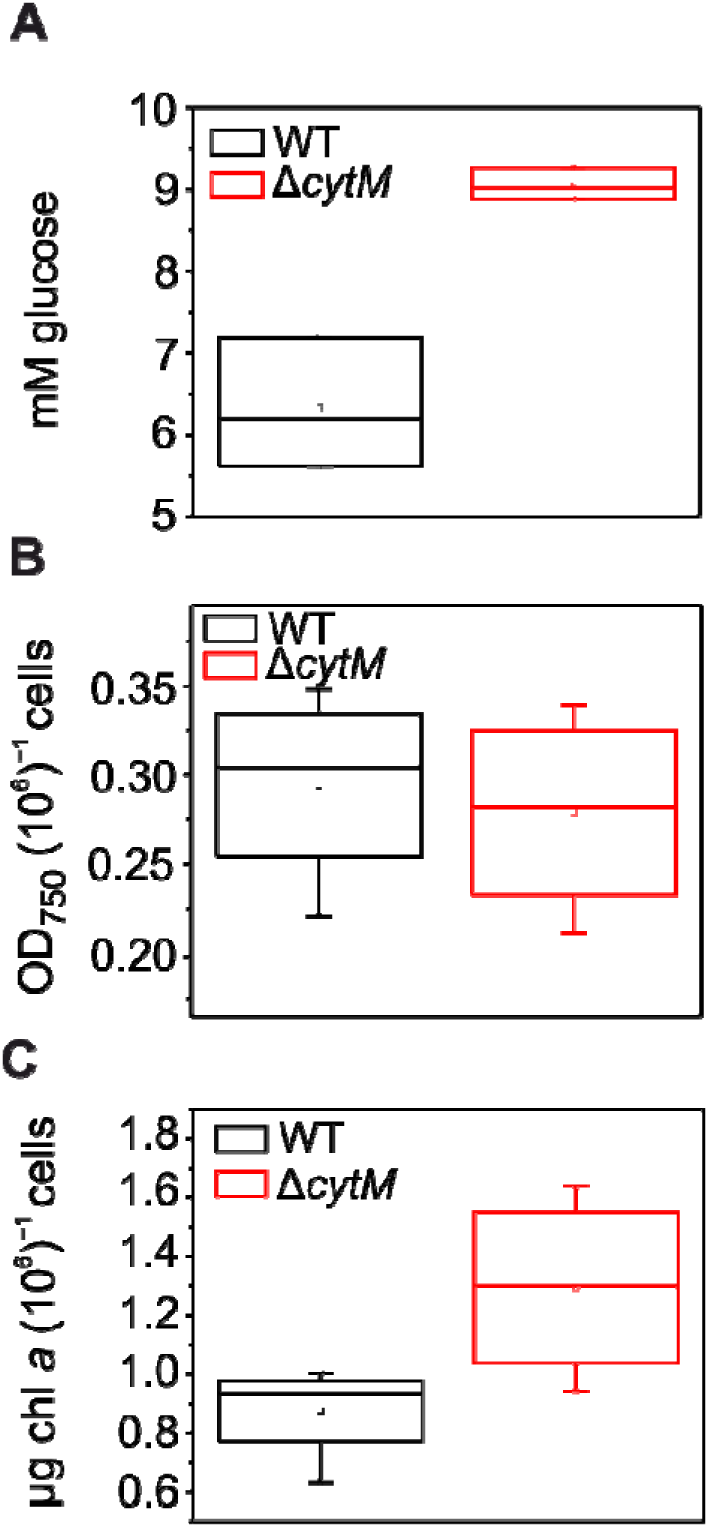
Glucose consumption, cellular chl content and cell number of WT and ΔCytM cultures on the third day of photomixotrophic growth. Amount of glucose consumed by the cells (A) was deduced from the remaining glucose in spent media. This number reflects the consumption of the whole culture rather than the glucose uptake rate of a given number of cells. Optical density per cell number (B) and cellular chl content (C) was determined. Values are means ± SD, n = three biological replicates. Cultures were grown photomixotrophically under constant 50 µmol photons m^−2^ s^−1^ illumination supplemented with 10 mM glucose. Samples were taken on the third day.

The effect of deleting *cytM* was tested then by culturing strains under photomixotrophic conditions but with different light regimes (Fig. 1C, D), either constant 10 µmol photons m^−2^ s^−1^ light (low light photomixotrophy) or 15 min 50 µmol photons m^−2^ s^−1^ light every 24 h (LAHG, light-activated heterotrophic growth). These cultures were supplemented with 10 mM starting glucose. Interestingly, under low light photomixotrophy, neither ΔCytM nor ΔCox/Cyd/CytM demonstrated a growth advantage compared to WT and ΔCox/Cyd, respectively. Under LAHG condition, ΔCytM grew faster than WT as previously reported (Hiraide et al 2015). The ΔCox/Cyd and ΔCox/Cyd/CytM mutants were unable to grow under LAHG. Previously it was reported that Cox is indispensable under this condition (Pils et al 1997).

We next examined the morphology of ΔCytM and WT cells on the third day of photomixotrophic growth (10 mM glucose, 50 µmol photons m^−2^ s^−1^ constant light), when the highest difference in OD_750_ was observed. The cell size, cell number per OD_750_ and chlorophyll (chl) concentration per cell was determined. No difference was observed in cell size between ΔCytM and WT (Supplemental Fig. S3), and the cell number per OD_750_ was similar in both strains (Fig. 2B), confirming that the difference in OD_750_ reflects higher growth. However, the chl *a* content per cell increased in ΔCytM (Fig. 2C), suggesting that the photosystem content or PSII/PSI ratio has been altered in this strain.

Overall, the most pronounced growth advantage of ΔCytM over the WT was observed when cells were exposed to a light intensity of 50 µmol photons m^−2^ s^−1^ and glucose concentration of 10 mM. Therefore, these conditions were used for all subsequent phenotyping experiments examining cells cultured photomixotrophically. The same phenotype manifested in the triple ΔCox/Cyd/CytM mutant, showing that Cox and Cyd are not required for the growth advantage. Moreover, we demonstrate that deletion of *cytM* leads to a higher cellular chl *a* content, which implies an altered photosynthetic machinery when cells are cultured photomixotrophically.

### Deletion of CytM circumvents the gradual over-reduction of the PQ-pool in photomixotrophy

To determine how deletion of CytM may alter the photosynthetic machinery, we first assessed the activity of PSII by probing chl fluorescence (Fig. 3) in WT and ΔCytM whole cells with multiple-turnover saturating pulses in dark, under far-red and under actinic red light. Compared to cells cultured photoautotrophically (Supplemental Fig. S4A), photomixotrophically grown WT cells demonstrated substantially higher initial fluorescence (F_0_) (Fig. 3C) and slower relaxation of pulse-induced fluorescence in the dark (see the F_m_^D^ relaxation on Fig. 3C). Interestingly, a considerable rise in steady-state fluorescence was observed under far-red light (Fig. 3C), despite the negligible actinic effect of far-red illumination on PSII. A similar rise in fluorescence was observed in photoautotrophically cultured WT when cells were measured in the presence of 3-(3,4-Dichlorophenyl)-1,1-dimethylurea (DCMU) (Supplemental Fig. S4C), a chemical, which occupies the Q_B_ site, thus blocking Q_A_-to-Q_B_ forward electron transfer in PSII. Most importantly, the F_s_ level under actinic light was considerably higher compared to cells grown photoautotrophically and firing saturating pulses barely increased fluorescence (see F_m_’ on Fig. 3C), implying a negligible effective PSII yield (Y(II)) (Supplemental Fig. S5A). Similar results were observed in a different WT *Synechocystis* substrain commonly used in our laboratory (Supplemental Fig. S6) and in cells exposed to longer periods of illumination (Supplemental Fig. S7A). Taken together, these results suggest a highly-reduced PQ-pool in photomixotrophically cultured WT under illumination.

**Figure 3.**
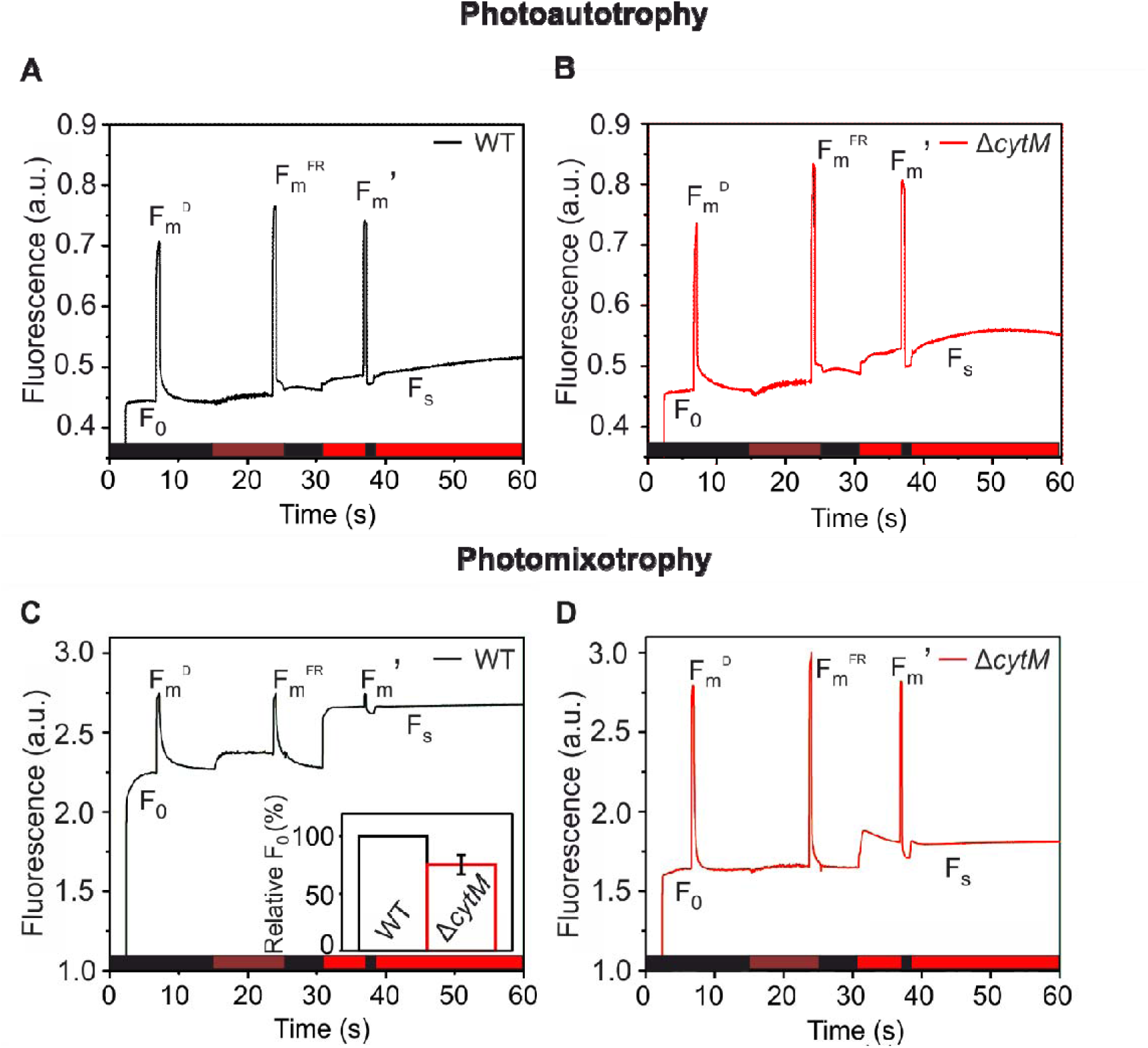
Fluorescence yield in WT and ΔCytM cells. Chl fluorescence of photoautotrophically (A, B) and photomixotrophically (C, D) grown WT and ΔCytM whole cells. Photoautotrophic and photomixotrophic cultures were grown under constant 50 µmol photons m^−2^ s^−1^ illumination for three days, with or without 10 mM glucose, respectively. Prior to measurements, the cell suspension was adjusted to 15 µg chl ml^−1^, resuspended in BG-11 supplemented with 10 mM glucose (C, D), and dark adapted for 15 min. Maximum fluorescence was determined by applying multiple turnover saturating pulse (500 ms, 5000 µmol photons m^−2^ s^−1^) in darkness (black bars), under 40 W m^−2^ far-red light (brown bars) and under 50 µmol photons m^−2^ s^−1^ actinic red light (red bars). F_0_, initial fluorescence; F_m_^D^, maximum fluorescence in dark; F_m_^FR^, maximum fluorescence in far-red; F_m_’, maximum fluorescence in actinic light; F_s_, steady state fluorescence in actinic light.

Compared to photomixotrophically grown WT, ΔCytM cultured under the same conditions demonstrated 24.8±8.3% lower F_0_ and the pulse-induced fluorescence relaxation in darkness was markedly faster (see F_m_^D^ on Fig. 3D). Far-red illumination did not increase fluorescence while saturating pulses greatly increased it (see F_m_’ on Fig. 4D), suggesting that the PSII effective yield Y(II) remained significantly higher, unlike in photomixotrophically grown WT cells (Supplemental Fig. S5A). Thus, in sharp contrast to WT, ΔCytM preserved a well-oxidized PQ-pool under photomixotrophy. Similarly, the triple mutant ΔCox/Cyd/CytM demonstrated high Y(II) compared to ΔCox/Cyd under photomixotrophy (Supplemental Fig. S7C-D).

**Figure 4.**
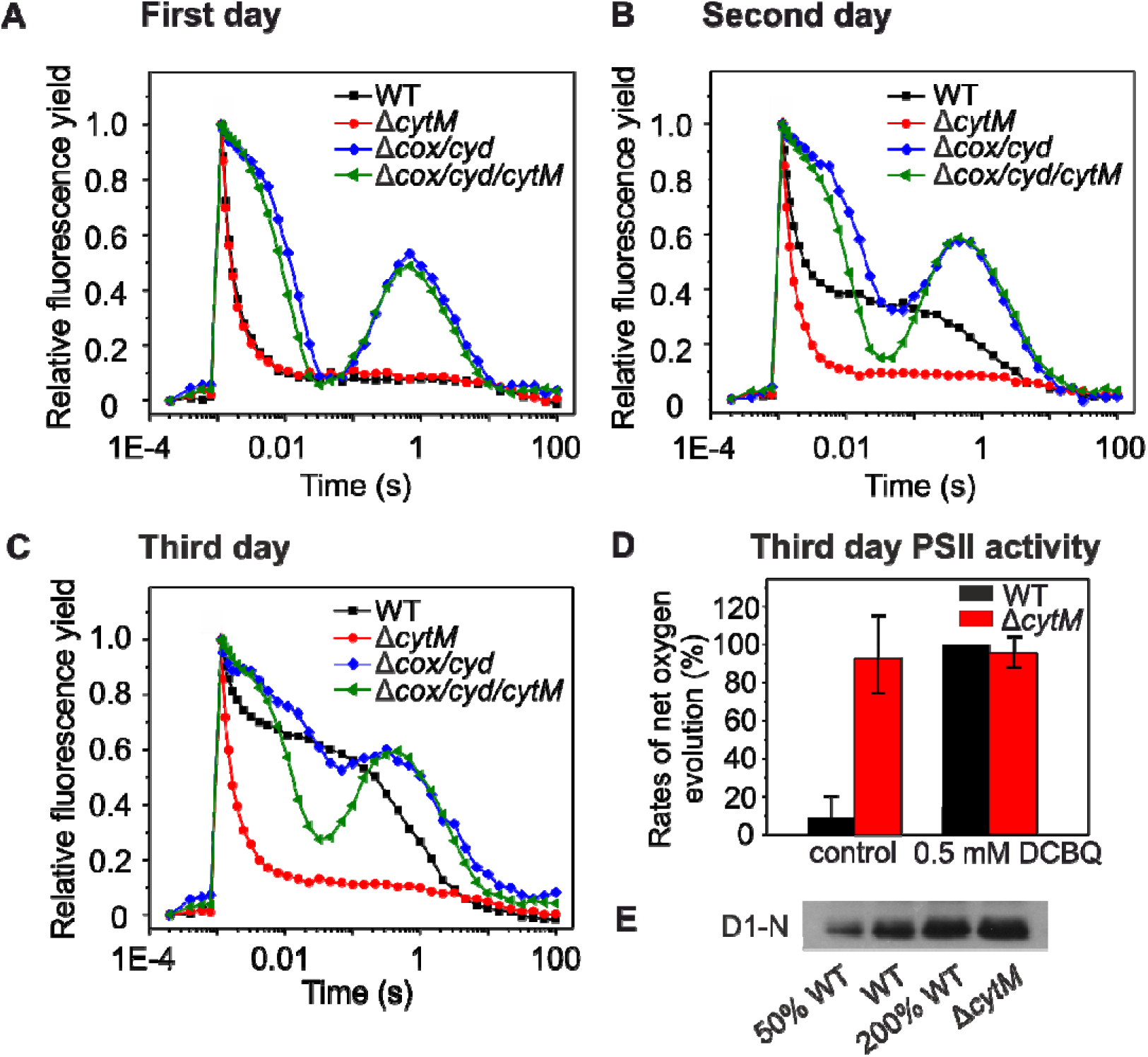
Relaxation of flash-induced fluorescence yield in cells exposed to darkness and quantification of O_2_ production capacity during photomixotrophic growth. Subsequent relaxation of fluorescence yields in the dark was measured after a single-turnover saturating pulse in photomixotrophically cultured cells taken on the first (A), second (B) and third day (C). Rates of net oxygen production (D) was determined in cells taken on the third day. O_2_ production was initiated with white light (1000 µmol photons m^−2^ s^−1^) in the absence (control) and in the presence of 0.5 mM DCBQ. Rates are expressed as µmol O_2_ mg chl^−1^ h^−1^, with DCBQ-treated WT considered as 100%. Values are means ± SD, n = three biological repeats. For numerically expressed rates of control samples, see Fig. 6. Immunoblot analysis with D1-N antibody (E) was performed on samples taken on the third day. 15 µg total protein extract was loaded per 100% lane, 50% and 200% corresponds to 7.5 µg and 30 µg, respectively.

To determine how over-reduction of the PQ-pool gradually increases in WT over three days of photomixotrophic growth, we monitored the redox kinetics of the PSII primary electron acceptor Q_A_ (Fig. 4) by firing a single-turnover saturating flash on dark-adapted cells then relaxation of the chl fluorescence yield was recorded in the period of subsequent darkness. No difference was observed between WT and ΔCytM cells cultured photoautotrophically (Supplemental Fig. S9) and on the first day of photomixotrophy, both WT and ΔCytM cells demonstrated typical flash-fluorescence relaxation in the darkness. On the second day, WT cells demonstrated a substantial slow-down in Q_A_-to-Q_B_ electron transfer reflected by slow decay kinetics (Fig. 4B), while on the third day, there was almost a complete loss of Q_A_-to-Q_B_ electron transfer (Fig. 4C). The kinetics from the third day resembled a curve recorded on photoautotrophically cultured WT supplemented with DCMU prior to the measurement (Supplemental Fig. S9). This supports the conclusion that Q_A_-to-Q_B_ electron transfer was almost completely inhibited in WT on the third day of photomixotrophy.

Interestingly, ΔCox/Cyd and ΔCox/Cyd/CytM displayed pronounced waving in the fluorescence yield relaxation kinetics (Fig. 4 A-C). The wave phenomenon is an unusual pattern in the decay of flash-induced chl fluorescence yield in the dark. The feature is characterized by a dip, corresponding to transient oxidation of Q_A_^-^ and a subsequent rise, reflecting re-reduction of the PQ-pool by NDH-1 (Deák et al 2014). During growth over the three-day period, the wave phenomenon in ΔCox/Cyd became less evident due to stronger inhibition of Q_A_-to-Q_B_ electron transfer. In contrast, ΔCox/Cyd/CytM displayed prominent waving during all three days of photomixotrophic growth, demonstrating that Q_A_^-^ re-oxidation was being sustained. Slight waving was reported previously (Ermakova et al 2016), although to our knowledge, this is the first study demonstrating that glucose induces a strong wave phenomenon in ΔCox/Cyd.

Next, we analysed net photosynthesis by probing the O_2_ production capacity of cells (Fig. 4D). When WT cells were grown photomixotrophically, only marginal net O_2_ production was observed on the third day. Strikingly, in the presence of the artificial electron acceptor, 2,6-dichloro-*p*-benzoquinone (DCBQ), the O_2_ evolving capacity of WT was restored. DCBQ accepts electrons from Q_A_ and/or Q_B_, virtually disconnecting PSII from the PQ-pool (Srivastava et al 1995). This suggests that PSII is fully functional in WT and that over-reduction of the PQ-pool hindered PSII electron transfer to PQ-pool. On the contrary, DCBQ did not increase O_2_ production in ΔCytM, implying that the PQ-pool is well oxidized. Immunoblotting performed on total protein extracts from WT and ΔCytM demonstrated a higher accumulation of PSII reaction center protein D1 in ΔCytM compared to WT (Fig 4E), suggesting that PSII is preserved in ΔCytM during the photomixotrophic growth.

These results demonstrate that during photomixotrophic growth, the PQ-pool gradually becomes over-reduced in WT, leading to drastically slower and eventually fully inhibited electron transfer from PSII to the PQ-pool on the third day. Deletion of CytM circumvents over-reduction of the PQ-pool and maintains PSII reaction center protein D1 amounts.

### ΔCytM has a larger pool of oxidizable PSI than WT in photomixotrophy

Next, we determined activity of PSI by monitoring the redox kinetics of P700, the primary electron donor of PSI (Fig. 5), which was performed simultaneously with chl fluorescence measurements (Fig. 3). First, the maximal amount of P700, P_m_, was determined (Fig. 5A). Compared to cells cultured under photoautotrophic conditions, WT cells grown photomixotrophically had 45.2±0.03% lower P_m_. Interestingly, the difference between ΔCytM cultured under photomixotrophic and photoautotrophic conditions was negligible (17.2±19.3%). Thus, under photomixotrophic conditions, ΔCytM had 132±18.7% higher maximum amounts of oxidizable P700 than WT (Fig. 5A). In line with this, immunoblotting revealed higher levels of a reaction center subunit of PSI, PsaB, in ΔCytM compared to WT under photomixotrophic growth (Fig. 5B). To determine the PSI:PSII ratio, samples were analysed at 77K by measuring chl fluorescence emission. No statistical difference was observed between WT and ΔCytM (Supplemental Fig. S10), demonstrating that the PSII:PSI ratio is the same in both strains.

**Figure 5.**
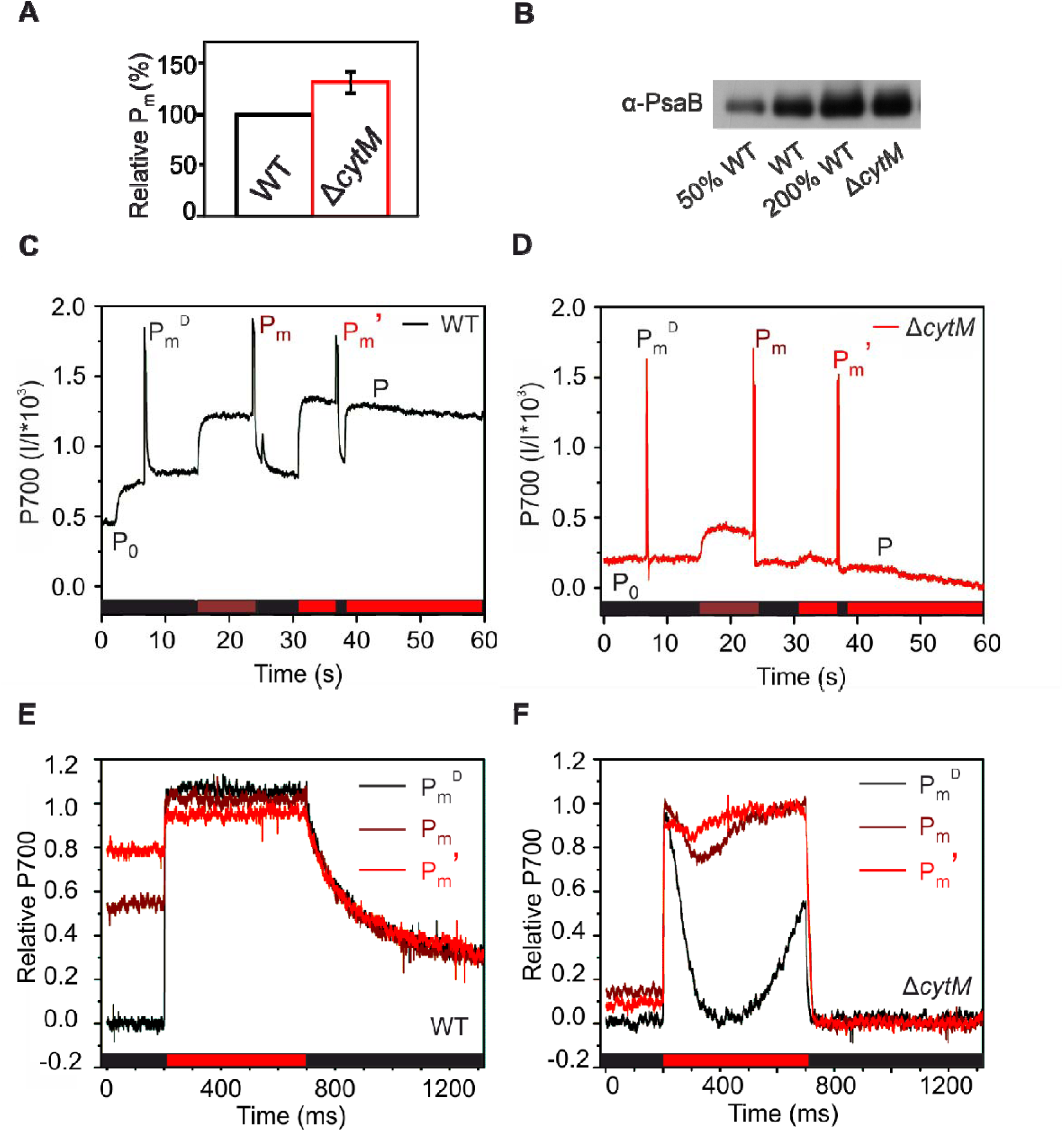
Characterization of PSI in cells cultured photomixotrophically. The maximal amount of oxidizable P700, P_m_ (A), and immunoblotting of PSI reaction center protein, PsaB (B), in cells cultured photomixotrophically. P700 oxidoreduction slow (C, D) and fast kinetics (E, F) was measured in parallel with fluorescence (Fig. 3). Fast kinetics curves (E, F) are normalized to P_m_ and referenced against their respective minimum P700 signal detected after the pulse. Cultivation, sample preparation and experimental parameters are similar to those detailed in Fig. 3. P_0_, initial P700; P_m_^D^, maximum P700 in darkness; P_m_^D^, maximum P700 under far-red light; P_m_’, maximum P700 under red actinic light.

The PSI effective yield Y(I), was also quantified, and was three times lower in photomixotrophically cultured WT cells compared to those grown photoautotrophically (Supplemental Fig S5B). This is due to a strong donor side limitation of PSI Y(ND), which demonstrates electron shortage to P700^+^ (Supplemental Fig S5C). In contrast, photomixotrophically cultured ΔCytM demonstrated similar Y(I) and only slightly increased Y(ND) compared to photoautotrophically cultured WT and ΔCytM (Supplemental Fig. S5B, C). As a result, ΔCytM had more than three times higher Y(I) than WT in photomixotrophy (Supplemental Fig. S5B).

Next, pulse-induced P700 fast kinetics were compared between photoautotrophically and photomixotrophically cultured WT (Fig. 5E) and ΔCytM (Fig 5F). These fast kinetics reveal the dynamics of P700 oxidoreduction during saturating pulses on millisecond scale. Saturating pulses are flashed in darkness (P_m_^D^) under far-red light (P_m_) and actinic red light (P_m_’). Typically, photoautotrophically cultured WT (Supplemental Fig. S11A) demonstrates transient P700^+^ re-reduction during light pulses. However, photomixotrophically grown WT did not exhibit the typical transient re-reduction (see P_m_^D^ on Fig. 5E). Importantly, P700^+^ relaxation after the pulse (Fig. 5E) was slower compared to that observed in photoautotrophically cultured cells (Supplemental Fig. S11A). Collectively, these results confirm that fewer electrons were transferred to P700^+^, leading to higher Y(ND) in photomixotrophically grown WT. Photomixotrophically cultured ΔCytM (Fig. 5F) displayed transient re-reduction during the pulses (see P_m_^D^, P_m_^FR^ and P_m_’ on Fig. 5F) and rapid relaxation after the pulse (Fig. 5F), resembling photoautotrophically cultured ΔCytM and WT (Supplemental Fig. S11A-B).

Here, we have shown that the effective yield of PSI in photomixotrophically cultured WT cells was considerably lower compared to photoautotrophically cultured cells, due to an electron shortage at P700^+^. This phenotype is eliminated by deleting *cytM*, as increased Y(I), higher amounts of oxidizable P700 (P_m_) and PsaB was observed in ΔCytM compared to WT on the third day of photomixotrophy.

### ΔCytM and ΔCox/Cyd/CytM sustain efficient net photosynthesis and CO_2_ fixation under photomixotrophy

To analyse real time gas exchange in photomixotrophically grown WT, ΔCytM, ΔCox/Cyd and ΔCox/Cyd/CytM (Fig. 6), O_2_ and CO_2_ fluxes in whole cells were monitored using MIMS (membrane inlet mass spectrometry) after enriching the samples with ^18^O_2_. In contrast to a classical oxygen electrode, which determines only net O_2_ changes, MIMS differentiates between gross photosynthetic O_2_ production by PSII and O_2_ consumption in light, mediated by flavodiiron proteins (Flv1-to-Flv4) and RTOs (Ermakova et al 2016, Santana-Sanchez et al 2019). Net O_2_ production is then calculated by subtracting the rates of O_2_ consumption in light from gross O_2_ production. Light-induced O_2_ consumption is calculated by subtracting the rates of O_2_ consumption in the dark from O_2_ consumption in the light.

**Figure 6.**
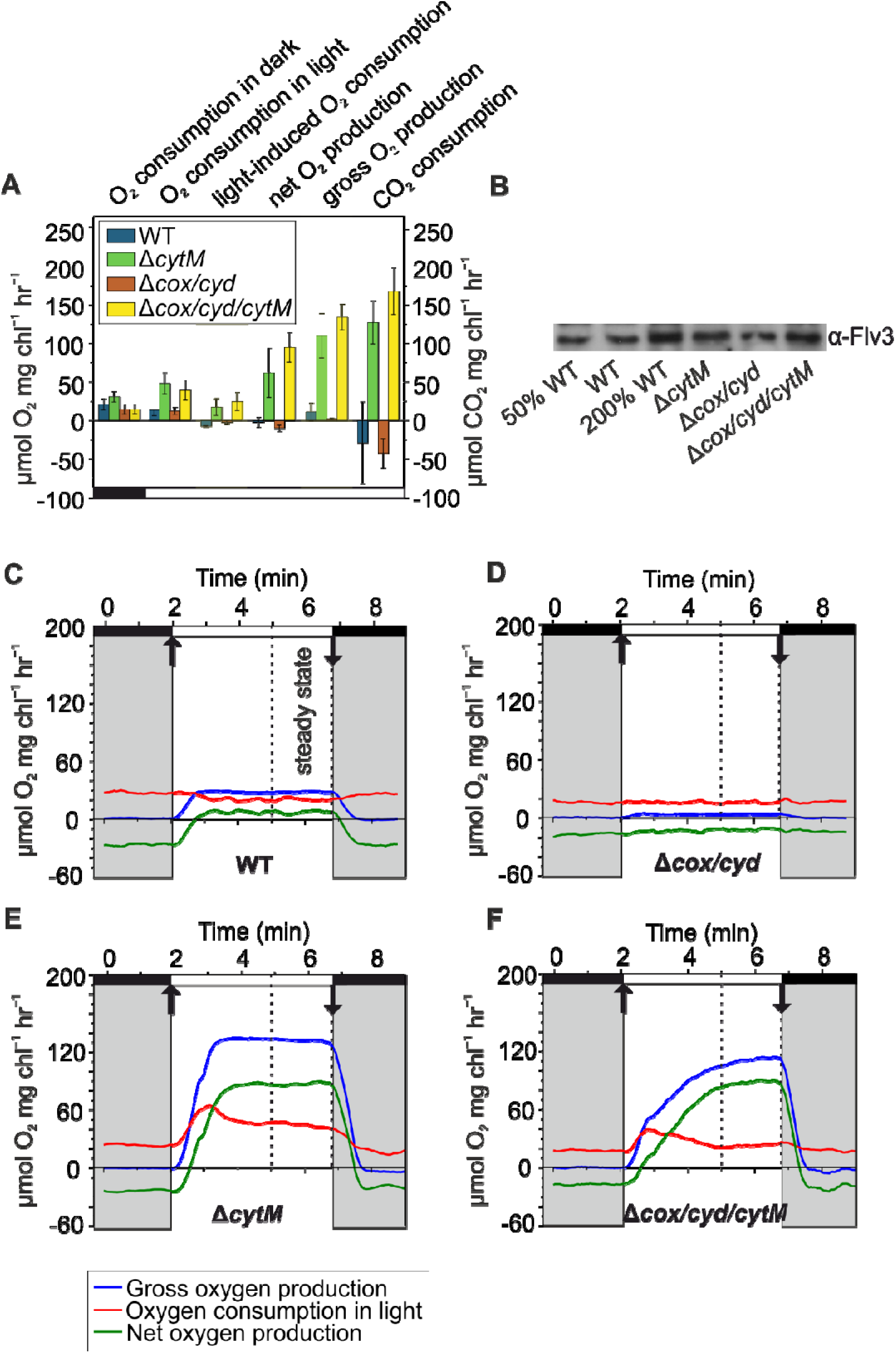
O_2_ and CO_2_ fluxes in photomixotrophically cultured WT, ΔCytM, ΔCox/Cyd and ΔCox/Cyd/CytM cells. Rates of O_2_ and CO_2_ fluxes in steady state (A). Values are means ± SD, n = 3-5 biological replicates. Analysis of total protein extracts by immmunoblotting with *α*-Flv3-specific antibody (B). 15 µg total protein was loaded per 100% lane, 50% and 200% corresponds to 7.5 µg and 30 µg, respectively. Kinetics of O_2_ flux rates in whole cells (C-F). Cultivation, sample preparation and experimental conditions are detailed in Fig. 3. In the light phase, 200 µmol photons m^−2^ s^−1^ constant white light was applied. Kinetics are representative of 3-6 biological replicates. The source data can be found in Supplemental table S2.

In WT under 200 µmol photons m^−2^ s^−1^ white light, O_2_ consumption and gross production rates were similar, resulting in nearly zero net photosynthetic O_2_ production. This is in line with the data obtained by O_2_ electrode (Fig. 4D). Corresponding to the minor net photosynthetic O_2_ production, the rate of CO_2_ consumption was negligible (Fig. 6A, Supplemental Fig. S12A). Importantly, no light-induced O_2_ consumption was observed in WT (Fig. 6A, C), although a substantial amount of Flv3 was detected by immunoblotting (Fig 6B). Although the thylakoid-localized RTOs, Cox and Cyd, were shown to be active in light (Ermakova et al 2016), a slight inhibition of respiratory O_2_ consumption under 200 µmol photons m^−2^ s^−1^ illumination occurred in WT. In contrast, ΔCytM exhibited a positive net O_2_ production rate and active CO_2_ consumption (Fig. 6E). Strikingly, gross O_2_ production was approximately 10 times higher compared to WT and ^18^O_2_ consumption in light followed a triphasic pattern, a characteristic trend reflecting the contribution of Flv1/3 and Flv2/4 to O_2_ consumption in light (Santana-Sanchez et al. 2019). The triphasic pattern in ΔCytM was observed as an initial burst of O_2_ consumption following the dark-to-light transition, which faded after 1-1.5 min and continued at a relatively constant rate (Fig. 6C). Accordingly, immunoblotting confirmed higher accumulation of the Flv3 proteins in ΔCytM. The rate of light-induced O_2_ consumption in ΔCytM is comparable to the reported values of photoautotrophically grown WT (Huokko et al 2017, Santana-Sanchez et al 2019). The dark respiration rate was higher in ΔCytM compared to WT, as previously observed when ΔCytM was cultured under dark, heterotrophic conditions (Hiraide et al 2015).

Similar to WT, ΔCox/Cyd (Fig 6C) showed barely any photosynthetic activity on the third day of photomixotrophic growth. During illumination, net O_2_ production remained negative, and CO_2_ consumption was found to be negligible (Fig. 6A, Supplemental Fig. S12B). Only residual gross O_2_ production was observed and O_2_ consumption was not stimulated by light (Fig 6A, D). Flv3 protein abundance in ΔCox/Cyd was comparable to WT (Fig. 6B). In sharp contrast to ΔCox/Cyd, ΔCox/Cyd/CytM demonstrated high PSII activity, producing O_2_ at a rate similar to ΔCytM. ΔCox/Cyd/CytM displayed a triphasic O_2_ consumption pattern under illumination (Fig. 6F) and the light-induced O_2_ consumption was comparable to that of ΔCytM in steady state (Fig. 6F). Compared to ΔCox/Cyd, ΔCox/Cyd/CytM had higher levels of Flv3 (Fig 6B). Notably, deleting *cytM* in the ΔCox/Cyd mutant did not enhance dark respiration, whereas ΔCytM had higher rates compared to WT.

To conclude, mutants lacking CytM sustained a steady electron flux towards O_2_ and CO_2_ in photomixotrophy, reflected by substantial net O_2_ production and active CO_2_ consumption under illumination both in ΔCytM and ΔCox/Cyd/CytM.

### Photomixotrophically cultured ΔCytM cells accumulate transport proteins and cofactor biosynthetic enzymes

In order to understand the metabolism of photomixotrophically grown WT and ΔCytM and how this affects the faster growth observed in the mutants, we analysed the total proteome by nLC-ESI-MS/MS via the data-dependent acquisition (DDA) method. Samples for analysis were collected on the second day, when both WT and ΔCytM cells were in late exponential phase and a significant growth difference was observed between the strains (Fig. 7A).

**Figure 7.**
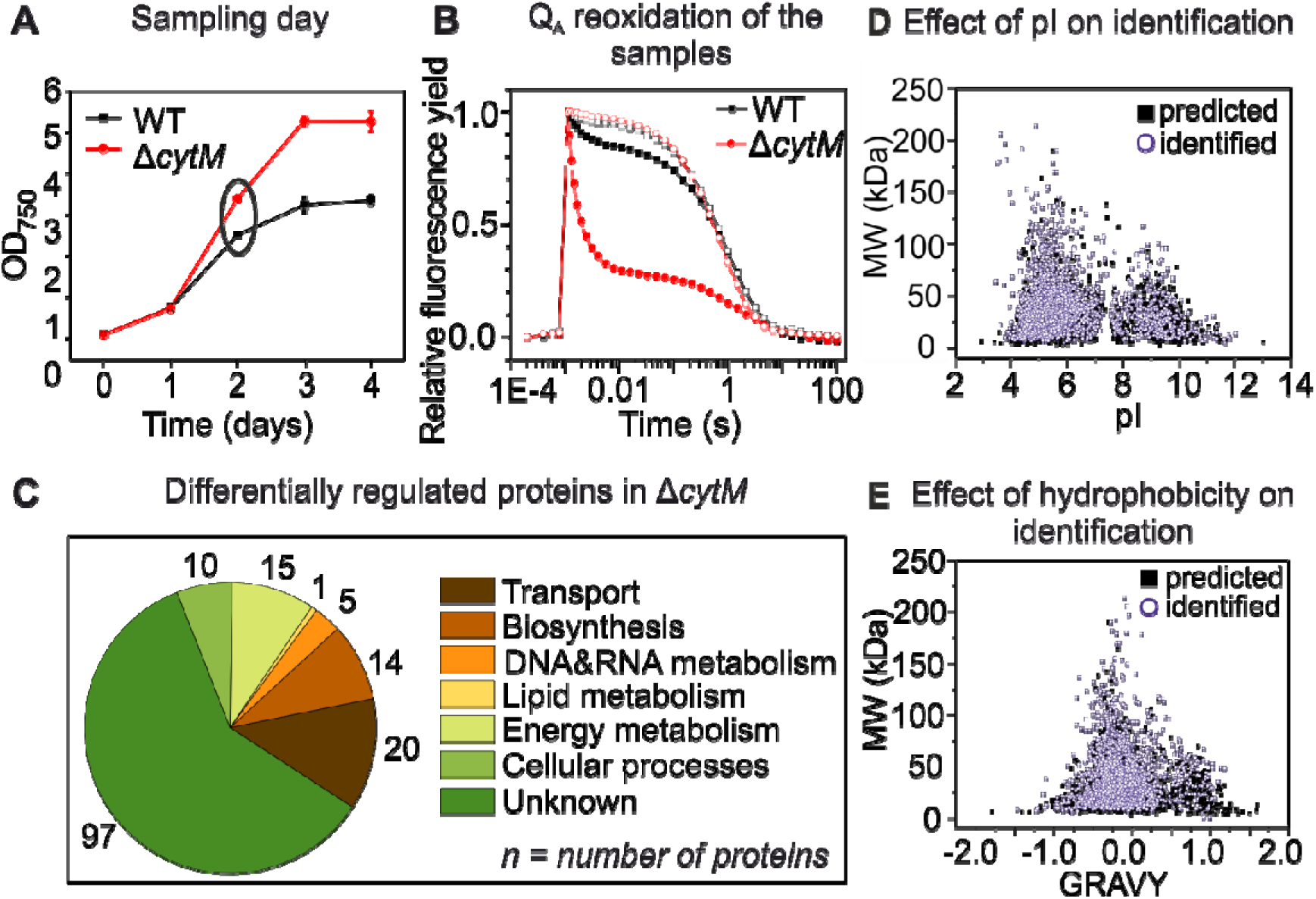
Characteristics at the sampling stage and functional classification of differentially regulated proteins in ΔCytM. Growth of the analysed cultures (A), with the ellipsis marking the sampling day. Cells were cultured similarly to those used in the biophysics analysis, except that the cells for proteomics were pre-cultivated under atmospheric CO_2_ in order to fully adapt the cells to these conditions. Importantly, extra pre-culturing step did not affect the growth of the experimental cultures. Relaxation of the flash-induced fluorescence yield in the dark (A) was measured in the absence (closed symbols) and in the presence of 20 µM DCMU (open symbols). Differentially regulated proteins in ΔCytM were grouped by functional classification (C). In total, 2415 proteins were identified, out of which 634 proteins were quantified and 162 were differentially regulated. The practical significance of differentially regulated proteins was set to FC >1.5 and FC <-1.5 (ANOVA p<0.05). Effect of isoelectric point (pI) (D) and hydrophobicity (GRAVY) (E) of the proteins on the identification rate was determined. Black squares mark all the 3507 predicted proteins in *Synechocystis*, lilac circles mark each proteins identified in WT and in ΔCytM.

In total, 2415 proteins were identified (Supplemental Table S3), despite the fact that the dataset was slightly biased against basic (Fig. 7D) and hydrophobic proteins (Fig. 7E), which is a known issue with this technique (Chandramouli and Qian 2009). Out of 2415 proteins, 634 were quantified, with 162 displaying a statistically different abundance in ΔCytM compared to WT (FC >1.5 and FC < −1.5 (ANOVA p<0.05) (Supplemental Table S4). The functional classification of differentially regulated proteins (Fig. 7C) revealed that apart from unknown or hypothetical proteins, mainly transporters and biosynthetic enzymes were altered in photomixotrophically cultured ΔCytM cells.

Table I shows a selection of proteins whose abundance was different in ΔCytM compared to WT. The highest fold change was observed in transport proteins. Among these, the constitutive low-affinity ABC-type phosphate transporters (PstA1, PstB1, PstB1’, PstC), periplasmic P_i_-binding proteins (SphX, PstS1) and extracellular lytic enzymes (PhoA, NucH) are more abundant in ΔCytM. Among proteins related to C_i_ uptake, a thylakoid *β*-type carbonic anhydrase, EcaB, was 2.32 times (P=7.50E-03) more abundant in ΔCytM. EcaB is a CupA/B-associated protein, proposed to regulate the activity of NDH-1_3_ (NDH-1 MS) and NDH-1_4_ (NDH-1 MS’) (Sun et al 2018). NDH-1_3_ facilitates inducible CO_2_-uptake, whereas NDH-1_4_ drives constitutive CO_2_-uptake (Ogawa et al 1991). CupB is exclusively found in the NDH-1_4_ complex and converts CO_2_ into HCO_3_^−^. Interestingly, no significant change was observed in the level of the glucose transporter GlcP, although the growth advantage of ΔCytM was glucose concentration-dependent.

**Table I.**
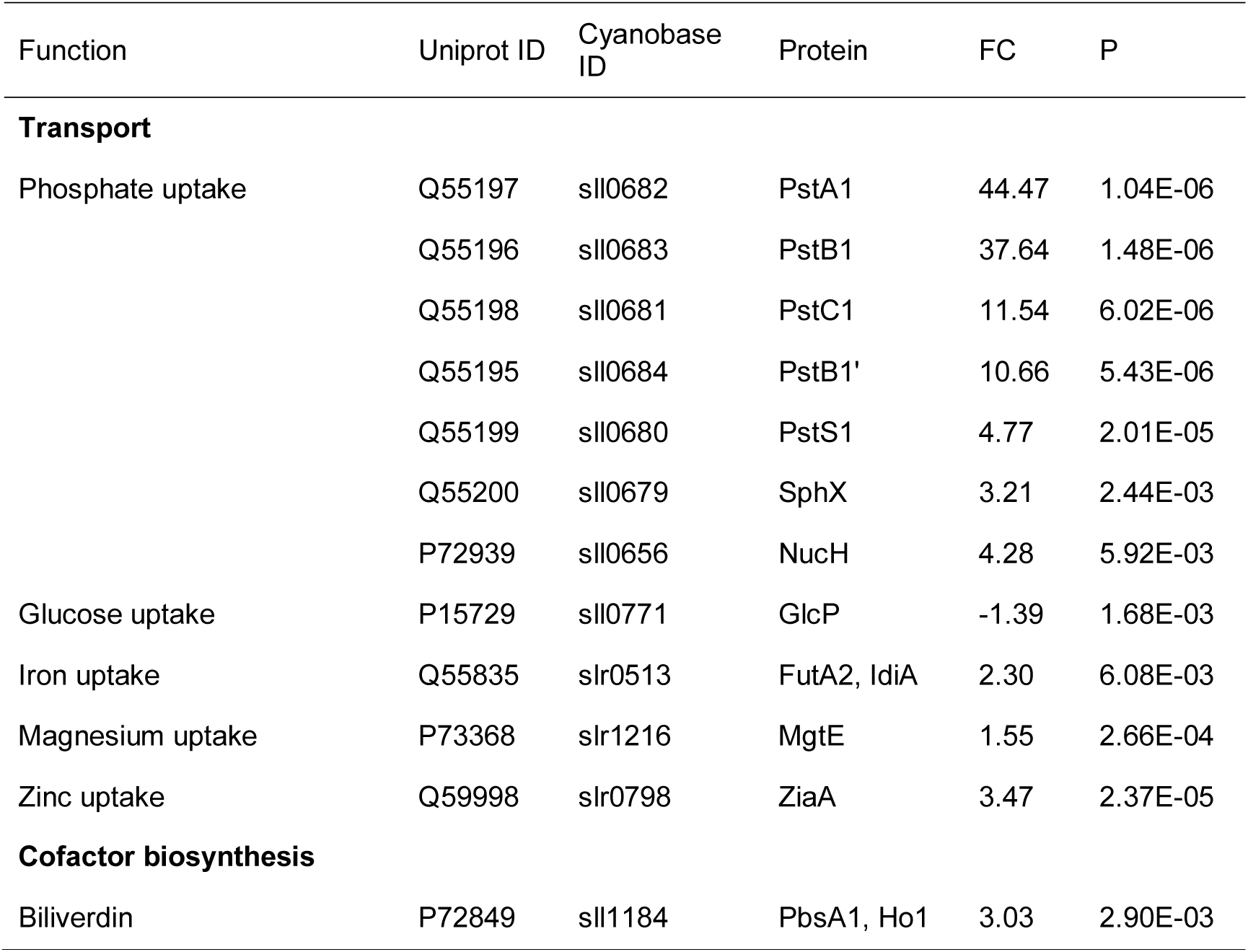
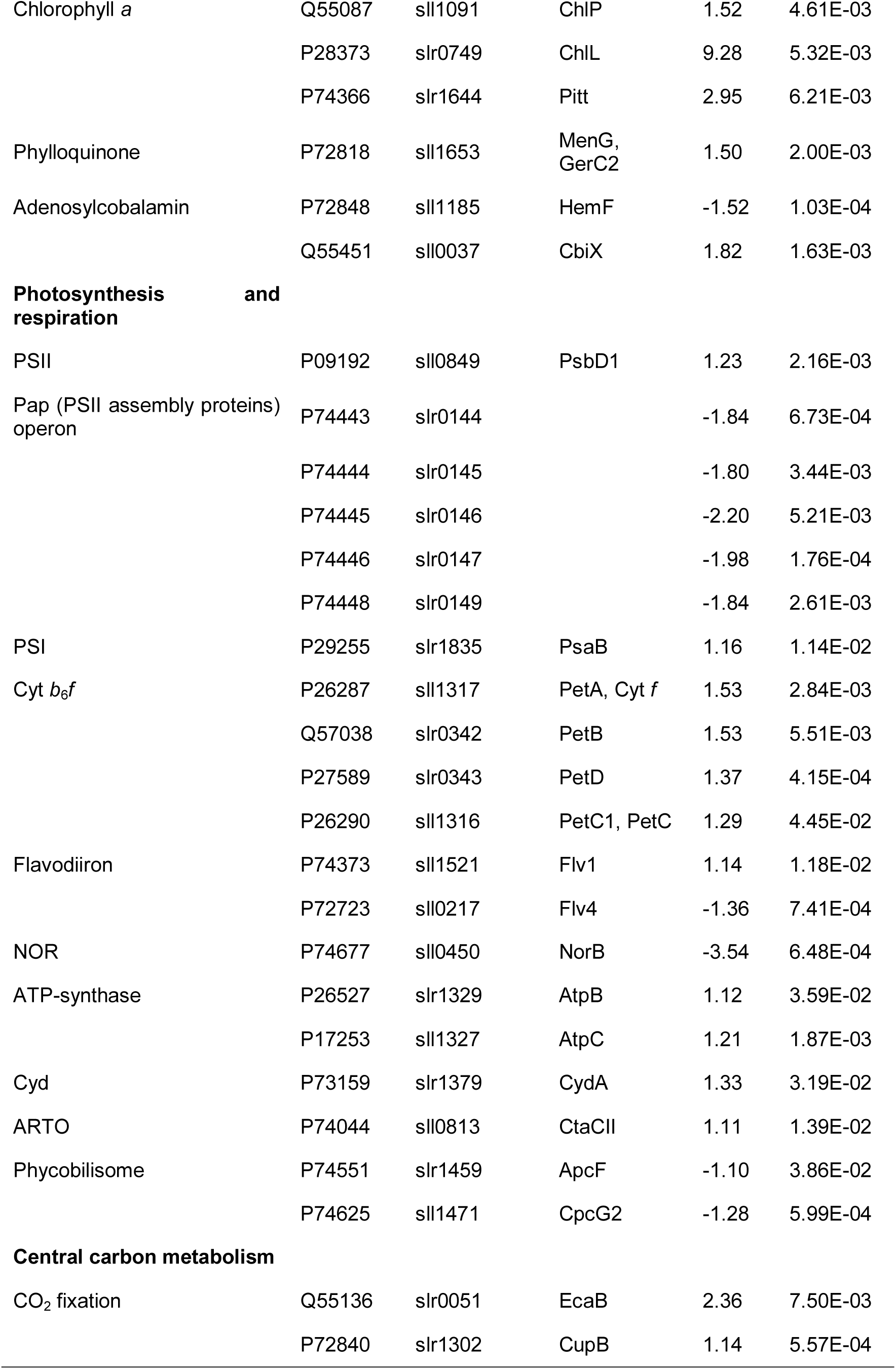
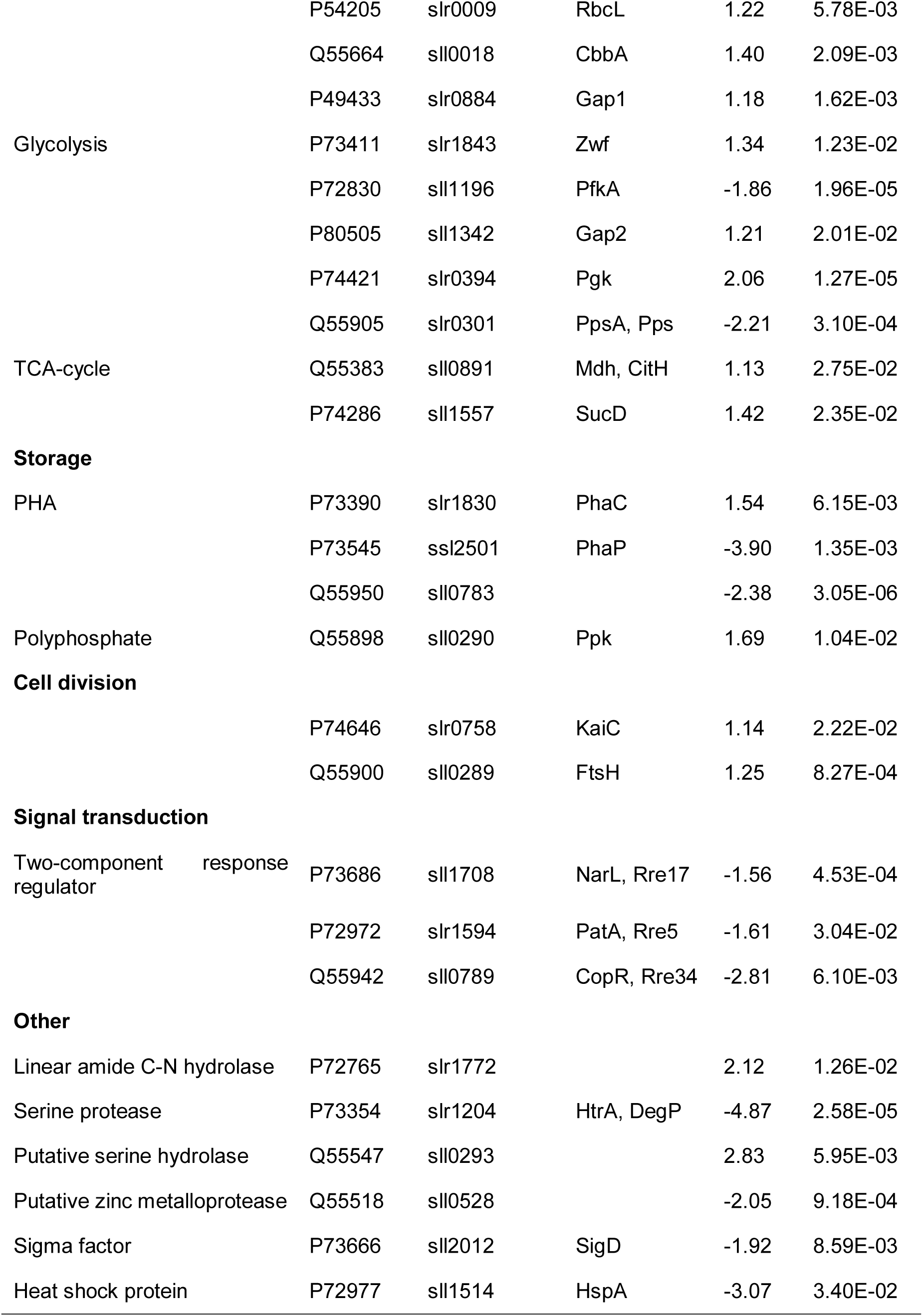
A selection of differentially expressed proteins in photomixotrophically cultured ΔCytM compared to WT. Cultivation and sampling are detailed in Fig. 7. Proteins were quantified by the DDA method and the practical significance of fold change was determined as FC≥1.5 and FC≤-1.5 (ANOVA p≤0.05). Significantly different proteins are marked with bold.

Chl *a* biosynthetic enzymes were found to accumulate in the mutant (Table I). ChlL, a subunit of the light-independent protochlorophyllide reductase (Wu and Vermaas 1995), and ChlP (4.61E-03), a geranylgeranyl reductase (Shpilyov *et al.* 2005), were 9.28 fold (P=5.32E-03) and 1.52 fold (P=4.61E-03) upregulated in ΔCytM, respectively. The incorporation of chl into photosystems likely increases due to the elevated level of Pitt, a protein contributing to the formation of photosynthetic pigments/proteins at the early stages of biogenesis (Schottkowski et al 2009). The ligand of the tetrapyrrole ring of chl is Mg^2+^ and accordingly, the magnesium uptake protein MgtE accumulated in ΔCytM along with a periplasmic iron-binding protein, FutA2, part of the complementary uptake-system of iron, a vital element of the photosynthetic machinery (Kranzler et al 2014). Among pigment biosynthetic enzymes, ΔCytM showed increased levels of the heme oxygenase Ho1, catalyzing the final step in the production of biliverdin (Willows et al 2000). Biliverdin is the precursor of phycocyanobilin, which is incorporated into phycobilisomes, the light-harvesting complexes of *Synechocystis*.

Among the photosynthetic proteins, the PSI reaction center subunit PsaB was found in equal amounts in WT and ΔCytM. However, immunoblotting with an anti-PsaB antibody demonstrated that ΔCytM contained higher amounts of PsaB than WT (Fig. 5B). This discrepancy may be due to the fact that despite the robustness of the MS-based DDA method, hydrophobic membrane proteins are prone to misquantification. Via MS analysis, quantification of *psbA* encoded D1 was not successful. Therefore, its abundance was only determined by immunoblotting (Fig. 4E), which revealed higher levels of D1 proteins in ΔCytM compared to WT. Interestingly and somewhat contradictorily, the amount of PSII assembly proteins encoded by the PAP-operon (Wegener et al 2008) decreased in the mutant. We also note that lower levels of NorB, a quinol-oxidizing nitric oxide reductase located in the plasma membrane (Büsch et al 2002), were observed in ΔCytM.

Since the growth advantage of ΔCytM is glucose-dependent, alterations are expected in the abundance of the intermediary carbon metabolic enzymes. In *Synechocystis*, roughly 100 enzymes participate in this metabolic network. In our study, 40 were quantified and surprisingly, only a few proteins were differentially regulated in ΔCytM. One notable example is phosphofructokinase PfkA, the key regulatory enzyme of the glycolytic Embden–Meyerhof–Parnas pathway, which was 1.86 times (P=1.96E-05) less abundant in ΔCytM, suggesting that carbon flux might be redirected into the Entner–Doudoroff or oxidative pentose phosphate pathways. Phosphoglycerate kinase Pgk, which is involved in each glycolytic pathway, was 2.06 times (P=1.27E-05) as abundant in ΔCytM. Phosphoenolpyruvate synthetase PpsA, a protein that catalyses the first step of gluconeogenesis, is 2.21 times (P=3.10E-04) less abundant in ΔCytM.

To conclude, global proteomic analysis revealed that photomixotrophically cultured ΔCytM accumulates transporter and chl biosynthetic proteins, while slight changes in the glycolytic and photosynthetic proteins were also observed.

## Discussion

The effect of importing and metabolising organic carbon on the bioenergetics properties of cyanobacteria over a long-term period is not fully understood. Previous studies have focused on the cellular changes following relatively short-term (from 10 min to 24 h) exposure to organic carbon (Lee et al 2007, Takahashi et al. 2008, Haimovich-Dayan et al 2011, Zilliges and Dau 2016). The majority of these reports suggest partial inhibition of photosynthetic activity, whereas some studies demonstrate increased net photosynthesis under air-level CO_2_ after 2 h exposure to 10 mM glucose (Haimovich-Dayan et al 2011). However, long-term changes to bioenergetics processes, particularly photosynthesis, remain to be elucidated. In this study, we investigated the effect of long-term photomixotrophic growth on WT and ΔCytM cells, most notably on the photosynthetic machinery, by analysing chlorophyll fluorescence, the redox kinetics of P700, real time O_2_ and CO_2_ fluxes and changes within the proteome.

### Gradual over-reduction of the PQ-pool limits photosynthesis in photomixotrophically cultured WT

By characterizing WT cells shifted from photoautotrophic to photomixotrophic conditions, we show that photosynthesis was markedly downscaled over three days of cultivation. This is deduced from the low PSII and PSI yield (Fig. 4C, D; Supplemental Fig. S5A, B) and most importantly, the negligible net O_2_ production (Fig. 6A, C) and CO_2_ fixation rates (Fig. 6A) on the third day. Since addition of an artificial PSII electron acceptor, DCBQ, restores O_2_ evolving activity of PSII (Fig. 5D), we can conclude that the highly reduced state of the PQ-pool limits electron flow from PSII and consequently, water splitting. However, residual PSII activity is ensured by circulating electrons in a water-water cycle. This was demonstrated by residual PSII gross O_2_ production (Fig. 6 A, C), nearly equalling O_2_ consumption in the light, resulting in practically zero net O_2_ production. Interestingly, PSI and FDPs fail to oxidize the over-reduced PQ-pool. High donor side limitation at PSI (Supplemental Fig. S5C) shows that the photosynthetic electrons accumulate in the PQ-pool and are not transferred to PSI. These results suggest that photosynthetic electron flow is restricted downstream to PSII.

Importantly, over-reduction of the PQ-pool increases gradually during photomixotrophic growth (Fig. 4). Based on thermoluminescence profiles, binding of quinones to PSII is delayed by glucose, although after two hours, the yield of PSII is not affected (Haimovich-Dayan et al 2011). We show here that after 24 hours growth (Fig. 4A), Q_A_^-^ reoxidation kinetics are comparable to photoautotrophically grown cells (Fig. S9), indicating that the PQ-pool is well oxidized. Over-reduction of the PQ pool occurs over the next two days when Q_A_-to-Q_B_ electron transfer becomes nearly completely blocked (Fig. 4B and 5C).

The gradual downscaling of photosynthesis could be due to a number of factors including: (i) spatial isolation of PSII via rearrangement in the thylakoid to another location or (ii) post-translational modifications of PSII. Rearrangement of thylakoid-localised complexes, specifically NDH-1 and SDH, has been observed in response to redox-regulated changes in the electron transport chain (Liu et al 2012). Applying the same analogy to PSII, the highly reduced state of the PQ-pool might trigger the complexes to arrange into a more sparse distribution during photomixotropic growth. Although cyanobacterial thylakoids are densely packed membranes (Kaňa et al 2013), lateral heterogeneity (Agarwal et al 2010) and mobility of PSII (Casella et al 2017) has been previously demonstrated. In the (ii) second scenario, post-translational modifications near the Q_B_-binding pocket hinders the effective oxidation of Q_A_^-^. D1 is phosphorylated at the C-terminus under certain conditions (Angeleri et al 2016), although the Q_B_-binding site resides at the N-terminus.

### Photomixotrophy does not alter photosynthetic electron transport in ΔCytM

Surprisingly, deletion of CytM reverses downscaling of photosynthesis in photomixotrophy, resulting in a profile similar to WT and ΔCytM cells grown under photoautotrophic conditions. Importantly, ΔCytM demonstrated unrestricted electron flow between PSII and PSI. The well oxidized PQ-pool allowed unrestricted Q_A_-to-Q_B_ electron transfer (Fig. 5C) and the rate of gross O_2_ production (Fig 6AE) was ten times higher in ΔCytM than it was in WT cells cultured under photomixotrophic conditions. Contrary to WT, ΔCytM did not demonstrate PSI donor-side limitation (Supplemental Fig. S5C). Finally, the abundance of D1 (Fig. 5E), PsaB (Fig.6B), and PetA and PetB (Table I), the core subunits of PSII, PSI and Cyt *b*_6_*f*, respectively, was higher in ΔCytM than in WT, although the PSI:PSII ratio was unaltered (Supplemental Fig. S10). As a consequence, the rate of net O_2_ production and CO_2_ consumption (Fig. 6-7) was significantly higher in ΔCytM, demonstrating that deletion of CytM conserves photosynthetic activity and circumvents over-reduction of the PQ-pool in photomixotrophy.

The exact mechanism by which ΔCytM alleviates blockage of the electron transport pathway was not elucidated in this work, nor has an exact role for this protein been determined in previous studies. CytM has been suggested to play a role in transferring electrons from Cyt *b*_6_*f* to Flv1/3, limiting productivity but providing a possible alternative route for safely transferring electrons to O_2_ (Hiraide et al 2015). However, given the low midpoint potential of CytM, a large energy barrier would have to be overcome in order for electron transfer downstream of Cyt *b*_6_*f* to occur (Cho et al 2000). Moreover, we demonstrated that the absence of CytM does not decrease O_2_ photoreduction driven by FDPs in ΔCox/Cyd/CytM (Fig. 6F), thus excluding this possibility. Recently, a cyanobacterial ferredoxin, Fed2, was shown to play a role in iron sensing and regulation of the IsiA antenna protein, a protein which is typically expressed when cells are exposed to low-iron conditions (Schorsch et al 2018). Similar to Fed2, it is possible that CytM plays a regulatory role in the cell, rather than being directly involved in electron transport.

Under conditions when cells are exposed to high concentrations of glucose or other sugars, CytM may regulate photosynthetic electron transport and carbon fixation by limiting CO_2_ uptake and decreasing the total amount of photosynthetic proteins, which in turn reduces photosynthesis. In line with this, we observed accumulation of EcaB in ΔCytM (Table I; Fig. 8). Enhanced EcaB levels likely results in greater inorganic carbon assimilation, higher carbon fixation, and increased turnover of NADPH, the terminal electron acceptor in linear photosynthetic electron transport. This in turn likely limits over-reduction of the photosynthetic electron transport chain.

**Fig. 8.**
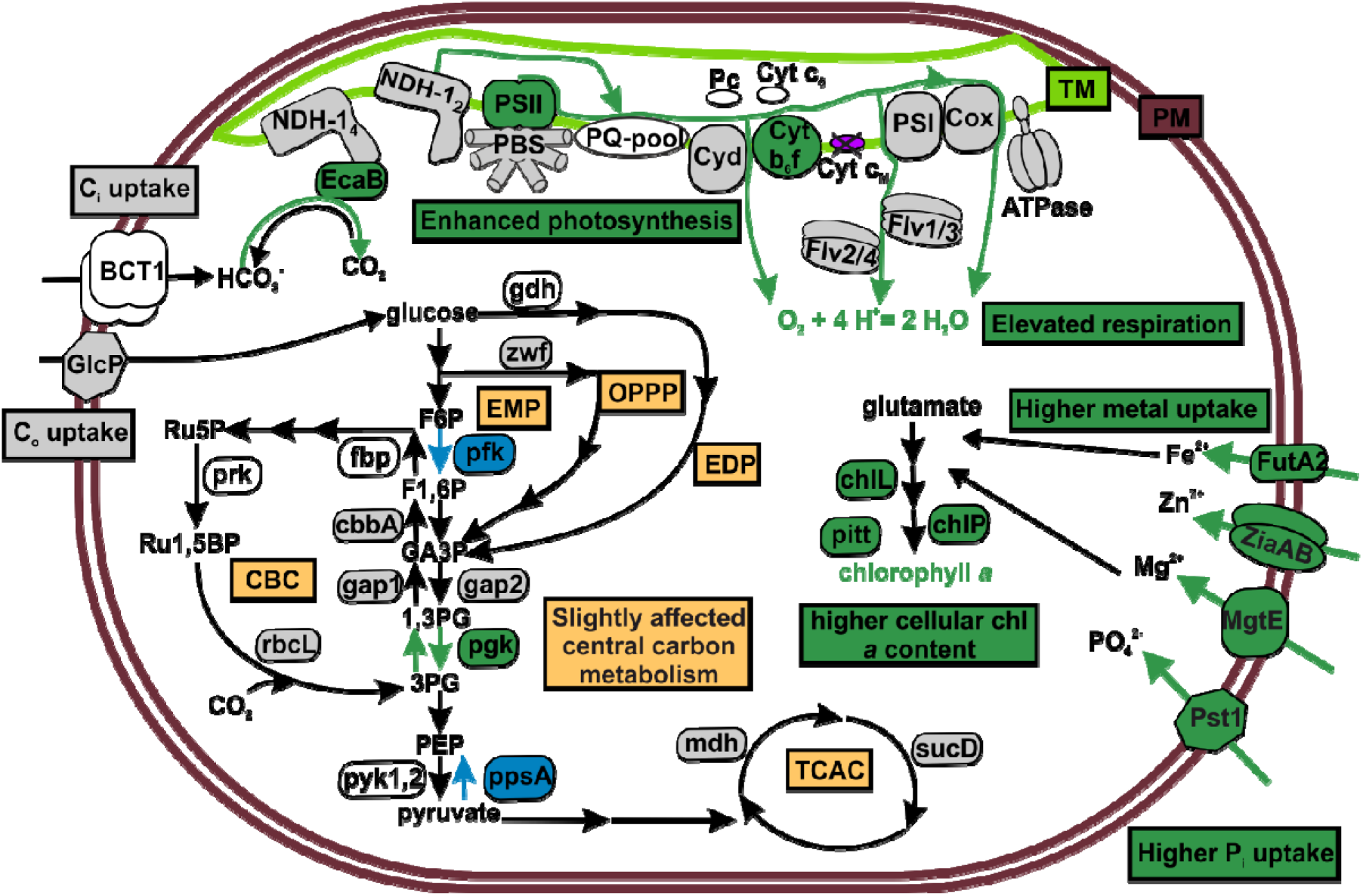
Schematic showing changes in the metabolism in photomixotrophically grown ΔCytM cells compared to WT. Proteins, compounds and metabolic routes with increased abundance or activity in ΔCytM relative to the WT are marked in green. Blue marks lower abundance in ΔCytM, grey marks unchanged and white marks undetermined abundance or activity. TM, thylakoid membrane; PM, plasma membrane.

Regardless of the exact role of CytM, it is clear that deletion of this protein significantly increases growth of *Synechocystis* in photomixotrophy (Fig. 1), in line with previous studies (Hiraide et al 2015). This is possibly due to an increase in photosynthetic capacity combined with efficient assimilation of glucose into central metabolism, resulting in greater biomass accumulation. This clearly results in increased production of proteins required for enhanced growth, including those involved in phosphate uptake (PstA1, PstB1, PstB1’, PstC) (Table I), import of Mg^2+^ (MgtE) Zn^2+^ (ZiaA) and Fe^2+^ (FutA2) and production of chl (ChlP, ChlL) (Table I; Fig. 8).

In conclusion, deletion of CytM allows *Synechocystis* to maintain efficient photosynthesis and enhanced growth under photomixotrophy. While we have not determined the exact function of CytM, we propose that it plays a role in reducing photosynthesis under natural conditions when both light intensity and glucose concentration fluctuates (Hieronymi and Macke et al 2010, Ittekkot et al 1985), and the redox state of the intertwined photosynthetic and respiratory electron transfer rapidly changes.

## MATERIALS AND METHODS

### Plasmid construction

The genome sequence of *Synechocystis* released 11.05.2004 was consulted via Cyanobase (http://genome.kazusa.or.jp/cyanobase) for primer design. Primers are listed in Supplemental Table S1. The *cytM* (*sll1245)* gene was deleted by amplifying a 906 bp fragment upstream of *cytM* using primers CytMleftfor and CytMleftrev and a 932 bp fragment downstream of *cytM* using primers CytMrightfor and CytMrightrev, followed by insertion of the respective fragments into the SacI/EcoR1 and XbaI/BamH1 sites of pUC19 to generate pCytM-1. The BamH1 digested npt1/sacRB cassette from pUM24Cm (Ried and Collmer, 1987) was inserted into the BamH1 site between the upstream and downstream fragments in pCytM-1 to generate pCytM-2.

### Construction of *cytM* deletion mutants

Unmarked mutants of *Synechocystis* lacking *cytM* were constructed via a two-step homologous recombination protocol according to Lea-Smith et al, 2016. To generate marked mutants approximately 1 µg of plasmid pCytM-2 was mixed with *Synechocystis* cells for 6 hours in liquid media, followed by incubation on BG-11 agar plates for approximately 24 hours. An additional 3 mL of agar containing kanamycin was added to the surface of the plate followed by further incubation for approximately 1-2 weeks. Transformants were subcultured to allow segregation of mutant alleles. Segregation was confirmed by PCR using primers CytMf and CytMr, which flank the deleted region. To remove the *npt*1/*sacRB* cassette to generate unmarked mutants, mutant lines were transformed with 1 µg of the markerless CytM-1 construct. Following incubation in BG-11 liquid media for 4 days and agar plates containing sucrose for a further 1-2 weeks, transformants were patched on kanamycin and sucrose plates. Sucrose resistant, kanamycin sensitive strains containing the unmarked deletion were confirmed by PCR using primers flanking the deleted region (Supplemental Fig. S2B). The ΔCox/Cyd/CytM unmarked strain was generated via the same method in the background of the unmarked ΔCox/Cyd strain (Lea-Smith et al., 2013).

### Cultivation

Pre-experimental cultures were grown in 30 ml BG-11 medium buffered with 10 mM TES-KOH (pH 8.2) in 100 ml Erlenmeyer flasks. Cultures were shaken at 120 rpm at 30°C and exposed to constant white fluorescent light of 50 µmol photons m^−2^ s^−1^ intensity in a Sanyo Environmental Test Chamber (Sanyo Co, Japan) which was saturated with 3% CO_2_. Starter cultures were inoculated at 0.1 OD_750_ and cultivated for three days with density typically reaching 2.5±0.5 OD_750_.

Experimental cultures for growth and photophysiological experiments were inoculated in 30 ml fresh BG-11 media at 0.1 OD_750_ from harvested pre-experimental cultures. The media was buffered with 10 mM TES-KOH (pH 8.2), the CO_2_ concentration was atmospheric, and cultures were agitated in 100 ml Erlenmeyer flasks at 120 rpm in AlgaeTRON AG130 cool-white LED chambers (PSI Instruments, Czech Republic). Growth was tested under constant light of 50 µmol photons m^−2^ with different glucose starting concentrations: (a) no glucose; (b) 5 mM glucose and (c) 10 mM glucose. At 10 mM glucose, additional light regimes were tested: (d) 10 µmol photons m^−2^ s^−1^ light and (e) 15 min 50 µmol photons m^−2^ s^−1^ every 24 h. For photophysiological studies, cells were cultivated under condition (c) for three days. For proteomics analysis, cells were cultivated similarly to (c), with the exception of an extra three day long pre-cultivation step at atmospheric CO_2_ without glucose.

### Cell counting, cell size determination

Cell number was determined with a Nexcelcom Cellometer X2 via the following method. Sample OD_750_ was adjusted to one, brightfield images were captured and the cell number was determined by the Nexcelcom software. In order to exclude the visual glitches falsely recognized as cells by the software, only the four most populous cell size group was averaged. Typically, three thousand cells were counted per plate.

### Glucose determination

Glucose concentration of the spent media was determined spectrophotometrically with the commercial High Sensitivity Glucose Assay Kit (Sigma-Aldrich, U.S.). Prior to measurements, the cell suspension was centrifuged at 5000 g for 10 min and the supernatant was filtered through a 0.2 µm filter.

### MIMS measurements

Gas fluxes of intact cells were measured using membrane inlet mass spectrometry. The in-house built system consists of a DW-1 oxygen electrode chamber (Hansatech Ltd., U.K.) connected to the vacuum line of a mass spectrometer (Prima PRO model, Thermo Scientific, U.S.). The sample was separated from the vacuum line by a Hansatech S4 PTFE membrane (Hansatech Ltd., U.K.). The chl *a* concentration of the sample was adjusted to 10 mg ml^−1^. Prior to measurements, the sample was enriched with 98 % ^18^O_2_ heavy isotope (CK Isotopes Limited, U.K.), the dissolved total carbon concentration was adjusted to 1.5 mM by adding NaHCO_3_, and then 10-15 min dark adaptation was applied. The measurement was performed in a semi-closed cuvette at 30°C with constant stirring. The light source was a 150 Watt, 21 V, EKE quartz halogen-powered fiber-optic illuminator (Fiber-Lite DC-950, Dolan-Jenner, U.S.). Rates were calculated as described previously (Beckmann et al 2009).

### Clark-type electrode measurements

PSII activity was tested in the presence of 0.5 mM 2,6-dichloro-*p*-benzoquinone (DCBQ) with a Clark-type oxygen electrode and chamber (Hansatech Ltd., U.K.). Cells were maintained at 30°C, with constant stirring, under 1000 µmol photons m^−2^ s^−1^ illumination using a Fiber-Lite DC-950 light source. Prior to the measurements, cells were resuspended in BG-11 (pH 8.2) supplemented with 10 mM glucose and the chl *a* concentration was adjusted to 7.5 µg ml^−1^. Rates of oxygen production was calculated using the Hansatech software.

### Chl fluorescence and P700 oxidoreduction measurements

Whole cell chl fluorescence was measured simultaneously with P700 with a pulse amplitude-modulated fluorometer (Dual-PAM-100, Walz, Germany). Prior to measurements, cells were resuspended in BG-11 (pH 8.2) supplemented with 10 mM glucose and the chl *a* concentration was adjusted to 15 µg ml^−1^. Measurements were performed at 30°C, and samples were initially incubated in darkness for 15 minutes with stirring. To determine P_m_, 30 s strong far-red light (720 nm, 40 W m^−2^) and red multiple turnover saturating pulses (MT) were applied. MT pulses were set to an intensity of 5000 µmol photons m^−2^ s^−1^ (width: 500 ms). Red (635 nm) actinic light was at an intensity of 50 µmol photons m^−2^ s^−1^ was used as background illumination. Photosynthetic parameters were calculated as described previously (Klughammer et al 2008 a,b).

Relaxation of flash-induced fluorescence yield was monitored using a fluorometer (FL3500, PSI Instruments, Czech Republic) outlined previously (Allahverdiyeva et al 2003). Prior to the measurement, cells were resuspended in BG-11 (pH 8.2) supplemented with 10 mM glucose, adjusted to 5 µg chl *a* ml^−1^ and dark adapted for 5 min. Curves were normalized to F_0_ and F_m_.

Fluorescence emission at 77K was determined using a USB4000-FL-450 fluorometer (Ocean Optics, USA). Prior to measurements, samples were adjusted to 7.5 mg chl ml^−1^ and light-adapted to identical conditions under which the cells were cultivated. Samples were excited at 440 nm. The curves were then normalized to their respective PSI emission peak at 723 nm.

### Western blotting

Total protein extraction, electrophoresis and immunoblotting was performed as described previously (Huokko et al 2019).

### MS analysis: sample preparation, data-dependent analysis, protein identification and quantitation

For data analysis, we used the proteome of *Synechocystis* sp. 6803 substr. Kazusa sequenced in 2004. Protein annotation was downloaded from Uniprot and Cyanobase. Hydrophobicity was determined via the GRAVY (grand average of hydropathy) index at www.gravy-calculator.de and pI was calculated via https://web.expasy.org/compute_pi/

Sample preparation for MS, data-dependent analysis and protein identification was performed as detailed previously (Huokko et al 2019). The mass spectrometry proteomics data was deposited to the ProteomeXchange Consortium via the PRIDE (Perez-Riverol et al 2019) partner repository with the dataset identifier PXD015246 and 10.6019/PXD015246.

## Supplemental data

**Supplemental figure S1.** Alignment of CytM from sequenced cyanobacterial species.

**Supplemental figure S2.** Generation of *cytM* deletion mutants in Synechocystis.

**Supplemental figure S3.** Cell size of WT and ΔCytM grown photomixotrophically and photoautotrophically.

**Supplemental figure S4**. Fluorescence transients of photoautotrophically cultivated WT and ΔCytM determined in the presence of 2 µM DCMU.

**Supplemental figure S5.** Photosynthetic parameters of WT and ΔCytM grown photomixotrophically and photoautotrophically.

**Supplemental Figure S6.** Fluorescence transients and P700 oxidoreduction on the third day of photomixotrophic growth of WT *Synechocystis* substrain.

**Supplemental Figure S7.** Fluorescence transients and P700 oxidoreduction kinetics of photomixotrophically grown WT, ΔCytM, ΔCox/Cyd and ΔCox/Cyd/CytM.

**Supplemental figure S8.** Fluorescence transients and P700 oxido-reduxtion kinetics of photoautotrophically grown WT, ΔCytM, ΔCox/Cyd and ΔCox/Cyd/CytM.

**Supplemental Figure S9.** Flash-induced increase of fluorescence yield and its relaxation in dark in photoautotrophically grown WT and ΔCytM.

**Supplemental Figure S10.** 77K steady state fluorescence emission spectra of WT and ΔCytM grown photomixotrophically.

**Supplemental Figure S11.** Fast kinetics of P700 oxidoreduction of WT and ΔCytM grown under photoautotrophic conditions.

**Supplemental Figure S12.** The rate of CO2 flux in photomixotrophically grown WT, ΔCytM, ΔCox/Cyd and ΔCox/Cyd/CytM.

**Supplemental Table S1.** List of oligonucleotides used in this study.

**Supplemental Table S2.** Rates of O_2_ and CO_2_ fluxes in photomixotrophically grown WT, ΔCytM, ΔCox/Cyd and ΔCox/Cyd/CytM.

**Supplemental Table S3.** Proteins identified by data-dependent analysis in photomixotrophically grown WT and ΔCytM.

**Supplemental Table S4.** Differentially expressed proteins in photomixotrophically grown ΔCytM versus WT.

## Acknowledgements

We thank Steffen Grebe for helping with Dual-PAM measurements and Ville Käpylä for laboratory assistance. MS analysis was performed at the Turku Proteomics Facility hosted by University of Turku and Åbo Akademi University, supported by Biocenter Finland. This work was supported by the Academy of Finland (project #315119 to Y.A. and the Finnish Center of Excellence, project #307335), the Nordforsk Nordic Center of Excellence ‘NordAqua’ (#82845) and the Waste Environmental Education Research Trust.

## Literature cited

Agarwal, R., Matros, A., Melzer, M., Mock, H.-P., Sainis, J.K., 2010. Heterogeneity in thylakoid membrane proteome of Synechocystis 6803. J. Proteomics 73, 976–991.

Allahverdiyeva, Y., Isojärvi, J., Zhang, P., Aro, E.-M., 2015. Cyanobacterial Oxygenic Photosynthesis is Protected by Flavodiiron Proteins. Life (Basel, Switzerland) 5, 716–43.

Allahverdiyeva, Y., Mustila, H., Ermakova, M., Bersanini, L., Richaud, P., Ajlani, G., Battchikova, N., Cournac, L., Aro, E.-M., 2013. Flavodiiron proteins Flv1 and Flv3 enable cyanobacterial growth and photosynthesis under fluctuating light. Proc. Natl. Acad. Sci. U. S. A. 110, 4111–6.

Angeleri, M., Muth-Pawlak, D., Aro, E.-M., Battchikova, N., 2016. Study of O - Phosphorylation Sites in Proteins Involved in Photosynthesis-Related Processes in Synechocystis sp. Strain PCC 6803: Application of the SRM Approach. J. Proteome Res. 15, 4638–4652.

Baers, L.L., Breckels, L.M., Mills, L.A., Gatto, L., Deery, M., Stevens, T.J., Howe, C.J., Lilley, K.S., Lea-Smith, D.J., 2019. Proteome mapping of a cyanobacterium reveals distinct compartment organisation and cell-dispersed metabolism. Plant Physiol. pp.00897.2019.

Beckmann, K., Messinger, J., Badger, M.R., Wydrzynski, T., Hillier, W., 2009. On-line mass spectrometry: membrane inlet sampling. Photosynth. Res. 102, 511–522.

Bernroitner, M., Tangl, D., Lucini, C., Furtmüller, P.G., Peschek, G.A., Obinger, C., 2009. Cyanobacterial cytochrome cM: Probing its role as electron donor for CuA of cytochrome c oxidase. Biochim. Biophys. Acta - Bioenerg. 1787, 135–143.

Bialek, W., Nelson, M., Tamiola, K., Kallas, T., Szczepaniak, A., 2008. Deeply branching c6-like cytochromes of cyanobacteria. Biochemistry 47, 5515–5522.

Bialek, W., Szczepaniak, A., Kolesinski, P., Kallas, T., 2016. Cryptic c 6-Like and c M Cytochromes of Cyanobacteria. In: Cytochrome Complexes: Evolution, Structures, Energy Transduction, and Signaling. Dordrecht: Springer, pp. 713–734.

Bothe, H., Schmitz, O., Yates, M.G., Newton, W.E., 2010. Nitrogen fixation and hydrogen metabolism in cyanobacteria. Microbiol. Mol. Biol. Rev. 74, 529–51.

Brown, K.A., Guo, Z., Tokmina-Lukaszewska, M., Scott, L.W., Lubner, C.E., Smolinski, S., Mulder, D.W., Bothner, B., King, P.W., 2019. The oxygen reduction reaction catalyzed by Synechocystis sp. PCC 6803 flavodiiron proteins. Sustain. Energy Fuels 3, 3191–3200.

Büsch, A., Friedrich, B., Cramm, R., 2002. Characterization of the norB gene, encoding nitric oxide reductase, in the nondenitrifying cyanobacterium Synechocystis sp. strain PCC6803. Appl. Environ. Microbiol. 68, 668–72.

Casella, S., Huang, F., Mason, D., Zhao, G.-Y., Johnson, G.N., Mullineaux, C.W., Liu, L.-N., 2017. Dissecting the Native Architecture and Dynamics of Cyanobacterial Photosynthetic Machinery. Mol. Plant 10, 1434–1448.

Chandramouli, K., Qian, P.-Y., 2009. Proteomics: challenges, techniques and possibilities to overcome biological sample complexity. Hum. Genomics Proteomics 2009.

Cho, Y.S., Pakrasi, H.B., Whitmarsh, J., 2000. Cytochrome cM from Synechocystis 6803. Eur. J. Biochem. 267, 1068–1074.

Deák, Z., Sass, L., Kiss, É., Vass, I., 2014. Characterization of wave phenomena in the relaxation of flash-induced chlorophyll fluorescence yield in cyanobacteria. Biochim. Biophys. Acta - Bioenerg. 1837, 1522–1532.

Durán, R. V, Hervás, M., De La Rosa, M.A., Navarro, J.A., 2004. The efficient functioning of photosynthesis and respiration in Synechocystis sp. PCC 6803 strictly requires the presence of either cytochrome c6 or plastocyanin. J. Biol. Chem. 279, 7229–33.

Ermakova, M., Huokko, T., Richaud, P., Bersanini, L., Howe, C.J., Lea-Smith, D.J., Peltier, G., Allahverdiyeva, Y., 2016. Distinguishing the Roles of Thylakoid Respiratory Terminal Oxidases in the Cyanobacterium Synechocystis sp. PCC 6803. Plant Physiol. 171, 1307–19.

Haimovich-Dayan, M., Kahlon, S., Hihara, Y., Hagemann, M., Ogawa, T., Ohad, I., Lieman-Hurwitz, J., Kaplan, A., 2011. Cross-talk between photomixotrophic growth and CO2-concentrating mechanism in Synechocystis sp. strain PCC 6803. Environ. Microbiol. 13, 1767–1777.

Heinz, S., Liauw, P., Nickelsen, J., Nowaczyk, M., 2016. Analysis of photosystem II biogenesis in cyanobacteria. Biochim. Biophys. Acta - Bioenerg. 1857, 274–287.

Helman, Y., Tchernov, D., Reinhold, L., Shibata, M., Ogawa, T., Schwarz, R., Ohad, I., Kaplan, A., 2003. Genes encoding A-type flavoproteins are essential for photoreduction of O2 in cyanobacteria. Curr. Biol. 13, 230–5.

Hieronymi, M., Macke, A., 2010. Spatiotemporal underwater light field fluctuations in the open ocean. J. Eur. Opt. Soc. Rapid Publ. 5, 10019s.

Hiraide, Y., Oshima, K., Fujisawa, T., Uesaka, K., Hirose, Y., Tsujimoto, R., Yamamoto, H., Okamoto, S., Nakamura, Y., Terauchi, K., Omata, T., Ihara, K., Hattori, M., Fujita, Y., 2015. Loss of cytochrome cM stimulates cyanobacterial heterotrophic growth in the dark. Plant Cell Physiol. 56, 334–45.

Huokko, T., Muth-Pawlak, D., Aro, E.-M., 2019. Thylakoid Localized Type 2 NAD(P)H Dehydrogenase NdbA Optimizes Light-Activated Heterotrophic Growth of Synechocystis sp. PCC 6803. Plant Cell Physiol. 60, 1386–1399.

Huokko, T., Muth-Pawlak, D., Battchikova, N., Allahverdiyeva, Y., Aro, E., 2017. Role of Type 2 NAD(P)H Dehydrogenase NdbC in Redox Regulation of Carbon Allocation in Synechocystis. Plant Physiol. 174, 1863–1880.

Ittekkot, V., Brockmann, U., Michaelis, W., Degens, E., 1981. Dissolved Free and Combined Carbohydrates During a Phytoplankton Bloom in the Northern North Sea. Mar. Ecol. Prog. Ser. 4, 299–305.

Kaňa, R., 2013. Mobility of photosynthetic proteins. Photosynth. Res. 116, 465–479.

Kanesaki, Y., Shiwa, Y., Tajima, N., Suzuki, M., Watanabe, S., Sato, N., Ikeuchi, M., Yoshikawa, H., 2012. Identification of substrain-specific mutations by massively parallel whole-genome resequencing of synechocystis sp. PCC 6803. DNA Res. 19, 67–79.

Kerfeld, C.A., Krogmann, D.W., 1998. Photosynthetic cytochromes c in cyanobacteria, algae and plants. Annu. Rev. Plant Physiol. Plant Mol. Biol. 49, 397–425.

Klughammer, C., Schreiber, U., 2008a. Complementary PS II quantum yields calculated from simple fluorescence parameters measured by PAM fluorometry and the Saturation Pulse method. PAM Appl. Notes 1, 27–35.

Klughammer, C., Schreiber, U., 2008b. Saturation Pulse method for assessment of energy conversion in PS I. PAM Appl. Notes 1, 11–14.

Kranzler, C., Lis, H., Finkel, O.M., Schmetterer, G., Shaked, Y., Keren, N., 2014. Coordinated transporter activity shapes high-affinity iron acquisition in cyanobacteria. ISME J. 8, 409–17.

Lea-Smith, David J., Bombelli, P., Vasudevan, R., Howe, C.J., 2016. Photosynthetic, respiratory and extracellular electron transport pathways in cyanobacteria. Biochim. Biophys. Acta - Bioenerg. 1857, 247–255.

Lea-Smith, D.J., Ross, N., Zori, M., Bendall, D.S., Dennis, J.S., Scott, S. a, Smith, A.G., Howe, C.J., 2013. Thylakoid terminal oxidases are essential for the cyanobacterium Synechocystis sp. PCC 6803 to survive rapidly changing light intensities. Plant Physiol. 162, 484–95.

Lea-Smith, David J, Vasudevan, R., Howe, C.J., 2016. Generation of Marked and Markerless Mutants in Model Cyanobacterial Species. J. Vis. Exp.

Lee, S., Ryu, J.-Y., Kim, S.Y., Jeon, J.-H., Song, J.Y., Cho, H.-T., Choi, S.-B., Choi, D., de Marsac, N.T., Park, Y.-I., 2007. Transcriptional regulation of the respiratory genes in the cyanobacterium Synechocystis sp. PCC 6803 during the early response to glucose feeding. Plant Physiol. 145, 1018–30.

Liu, L.-N., Bryan, S.J., Huang, F., Yu, J., Nixon, P.J., Rich, P.R., Mullineaux, C.W., 2012. Control of electron transport routes through redox-regulated redistribution of respiratory complexes. Proc. Natl. Acad. Sci. 109, 11431–11436.

Malakhov, M.P., Malakhova, O.A., Murata, N., 1999. Balanced regulation of expression of the gene for cytochrome cM and that of genes for plastocyanin and cytochrome c6 in Synechocystis. FEBS Lett. 444, 281–284.

Malakhov, M.P., Wada, H., Los, D.A., Semenenko, V.E., Murata, N., 1994. A New Type of Cytochrome c from Synechocystis PCC6803. J. Plant Physiol. 144, 259–264.

Manna, P., Vermaas, W., 1997. Lumenal proteins involved in respiratory electron transport in the cyanobacterium Synechocystis sp. PCC6803. Plant Mol. Biol. 35, 407–416.

Molina-Heredia, F.P., Balme, A., Hervás, M., Navarro, J.A., De la Rosa, M.A., 2002. A comparative structural and functional analysis of cytochrome c M, cytochrome c 6 and plastocyanin from the cyanobacterium Synechocystis sp. PCC 6803. FEBS Lett. 517, 50–54.

Moore, L.R., 2013. More mixotrophy in the marine microbial mix. Proc. Natl. Acad. Sci. 110, 8323–8324.

Mullineaux, C.W., 2014. Co-existence of photosynthetic and respiratory activities in cyanobacterial thylakoid membranes. Biochim. Biophys. Acta - Bioenerg. 1837, 503–511.

Mustila, H., Paananen, P., Battchikova, N., Santana-Sánchez, A., Muth-Pawlak, D., Hagemann, M., Aro, E.-M., Allahverdiyeva, Y., 2016. The Flavodiiron Protein Flv3 Functions as a Homo-Oligomer During Stress Acclimation and is Distinct from the Flv1/Flv3 Hetero-Oligomer Specific to the O2 Photoreduction Pathway. Plant Cell Physiol.

Ogawa, T., 1991. A gene homologous to the subunit-2 gene of NADH dehydrogenase is essential to inorganic carbon transport of Synechocystis PCC6803. Proc. Natl. Acad. Sci. 88, 4275–4279.

Perez-Riverol, Y., Csordas, A., Bai, J., Bernal-Llinares, M., Hewapathirana, S., Kundu, D.J., Inuganti, A., Griss, J., Mayer, G., Eisenacher, M., Pérez, E., Uszkoreit, J., Pfeuffer, J., Sachsenberg, T., Yilmaz, S., Tiwary, S., Cox, J., Audain, E., Walzer, M., Jarnuczak, A.F., Ternent, T., Brazma, A., Vizcaíno, J.A., 2019. The PRIDE database and related tools and resources in 2019: improving support for quantification data. Nucleic Acids Res. 47, D442–D450.

Pils, D., 1997. Evidence for in vivo activity of three distinct respiratory terminal oxidases in the cyanobacterium Synechocystis sp. strain PCC6803. FEMS Microbiol. Lett. 152, 83–88.

Ried, J.L., Collmer, A., 1987. An nptI-sacB-sacR cartridge for constructing directed, unmarked mutations in gram-negative bacteria by marker exchange-eviction mutagenesis. Gene 57, 239–46.

Santana-Sanchez, A., Solymosi, D., Mustila, H., Bersanini, L., Aro, E.-M., Allahverdiyeva, Y., 2019. Flavodiiron proteins 1–to-4 function in versatile combinations in O2 photoreduction in cyanobacteria. Elife 8.

Schorsch, M., Kramer, M., Goss, T., Eisenhut, M., Robinson, N., Osman, D., Wilde, A., Sadaf, S., Brückler, H., Walder, L., Scheibe, R., Hase, T., Hanke, G.T., 2018. A unique ferredoxin acts as a player in the low-iron response of photosynthetic organisms. Proc. Natl. Acad. Sci. U. S. A. 115, E12111–E12120.

Schottkowski, M., Ratke, J., Oster, U., Nowaczyk, M., Nickelsen, J., 2009. Pitt, a Novel Tetratricopeptide Repeat Protein Involved in Light-Dependent Chlorophyll Biosynthesis and Thylakoid Membrane Biogenesis in Synechocystis sp. PCC 6803. Mol. Plant 2, 1289–1297.

Schuller, J.M., Birrell, J.A., Tanaka, H., Konuma, T., Wulfhorst, H., Cox, N., Schuller, S.K., Thiemann, J., Lubitz, W., Sétif, P., Ikegami, T., Engel, B.D., Kurisu, G., Nowaczyk, M.M., 2019. Structural adaptations of photosynthetic complex I enable ferredoxin-dependent electron transfer. Science (80-.). 363, 257–260.

Schultze, M., Forberich, B., Rexroth, S., Dyczmons, N.G., Roegner, M., Appel, J., 2009. Localization of cytochrome b6f complexes implies an incomplete respiratory chain in cytoplasmic membranes of the cyanobacterium Synechocystis sp. PCC 6803. Biochim. Biophys. Acta - Bioenerg. 1787, 1479–1485.

Shen, J.R., Inoue, Y., 1993. Binding and functional properties of two new extrinsic components, cytochrome c-550 and a 12-kDa protein, in cyanobacterial photosystem II. Biochemistry 32, 1825–1832.

Shpilyov, A. V., Zinchenko, V. V., Shestakov, S. V., Grimm, B., Lokstein, H., 2005. Inactivation of the geranylgeranyl reductase (ChlP) gene in the cyanobacterium Synechocystis sp. PCC 6803. Biochim. Biophys. Acta - Bioenerg. 1706, 195–203.

Smith, A.J., 1983. Modes of cyanobacterial carbon metabolism. Ann. l’Institut Pasteur / Microbiol. 134, 93–113.

Sonoda, M., Kitano, K., Katoh, A., Katoh, H., Ohkawa, H., Ogawa, T., 1997. Size of cotA and identification of the gene product in Synechocystis sp. strain PCC6803. J. Bacteriol. 179, 3845–3850.

Srivastava, A., Strasser, R.J., Govindjee, 1995. Differential effects of dimethylbenzoquinone and dichlorobenzoquinone on chlorophyll fluorescence transient in spinach thylakoids. J. Photochem. Photobiol. B Biol. 31, 163–169.

Stal, L.J., Moezelaar, R., 1997. Fermentation in cyanobacteria. FEMS Microbiol. Rev. 21, 179–211.

Sun, N., Han, X., Xu, M., Kaplan, A., Espie, G.S., Mi, H., 2018. A thylakoid-located carbonic anhydrase regulates CO 2 uptake in the cyanobacterium Synechocystis sp. PCC 6803. New Phytol. nph. 15575.

Takahashi, H., Uchimiya, H., Hihara, Y., 2008. Difference in metabolite levels between photoautotrophic and photomixotrophic cultures of Synechocystis sp. PCC 6803 examined by capillary electrophoresis electrospray ionization mass spectrometry. J. Exp. Bot. 59, 3009–18.

Teeling, H., Fuchs, B.M., Becher, D., Klockow, C., Gardebrecht, A., Bennke, C.M., Kassabgy, M., Huang, S., Mann, A.J., Waldmann, J., Weber, M., Klindworth, A., Otto, A., Lange, J., Bernhardt, J., Reinsch, C., Hecker, M., Peplies, J., Bockelmann, F.D., Callies, U., Gerdts, G., Wichels, A., Wiltshire, K.H., Glöckner, F.O., Schweder, T., Amann, R., 2012. Substrate-controlled succession of marine bacterioplankton populations induced by a phytoplankton bloom. Science 336, 608–611.

Vermaas, W.F.J., 2001. Photosynthesis and respiration in cyanobacteria. In: Encyclopedia of Life Sciences. pp. 1–7.

Vicente, J.B., Gomes, C.M., Wasserfallen, A., Teixeira, M., 2002. Module fusion in an A-type flavoprotein from the cyanobacterium Synechocystis condenses a multiple-component pathway in a single polypeptide chain. Biochem. Biophys. Res. Commun. 294, 82–87.

Wegener, K.M., Welsh, E.A., Thornton, L.E., Keren, N., Jacobs, J.M., Hixson, K.K., Monroe, M.E., Camp, D.G., Smith, R.D., Pakrasi, H.B., 2008. High sensitivity proteomics assisted discovery of a novel operon involved in the assembly of photosystem II, a membrane protein complex. J. Biol. Chem. 283, 27829–37.

Willows, R.D., Mayer, S.M., Foulk, M.S., DeLong, A., Hanson, K., Chory, J., Beale, S.I., 2000. Phytobilin biosynthesis: the Synechocystis sp. PCC 6803 heme oxygenase-encoding ho1 gene complements a phytochrome-deficient Arabidopsis thaliana hy1 mutant. Plant Mol. Biol. 43, 113–120.

Wu, Q., Vermaas, W.F., 1995. Light-dependent chlorophyll a biosynthesis upon chlL deletion in wild-type and photosystem I-less strains of the cyanobacterium Synechocystis sp. PCC 6803. Plant Mol. Biol. 29, 933–45.

Zilliges, Y., Dau, H., 2016. Unexpected capacity for organic carbon assimilation by Thermosynechococcus elongatus, a crucial photosynthetic model organism. FEBS Lett. 590, 962–970.

Zubkov, M. V., Tarran, G.A., 2008. High bacterivory by the smallest phytoplankton in the North Atlantic Ocean. Nature 455, 224–226.

