## Supplementary figures and images for "Cytochrome c_M_ downscales photosynthesis under photomixotrophy in *Synechocystis* sp. PCC 6803"

### Suppl data

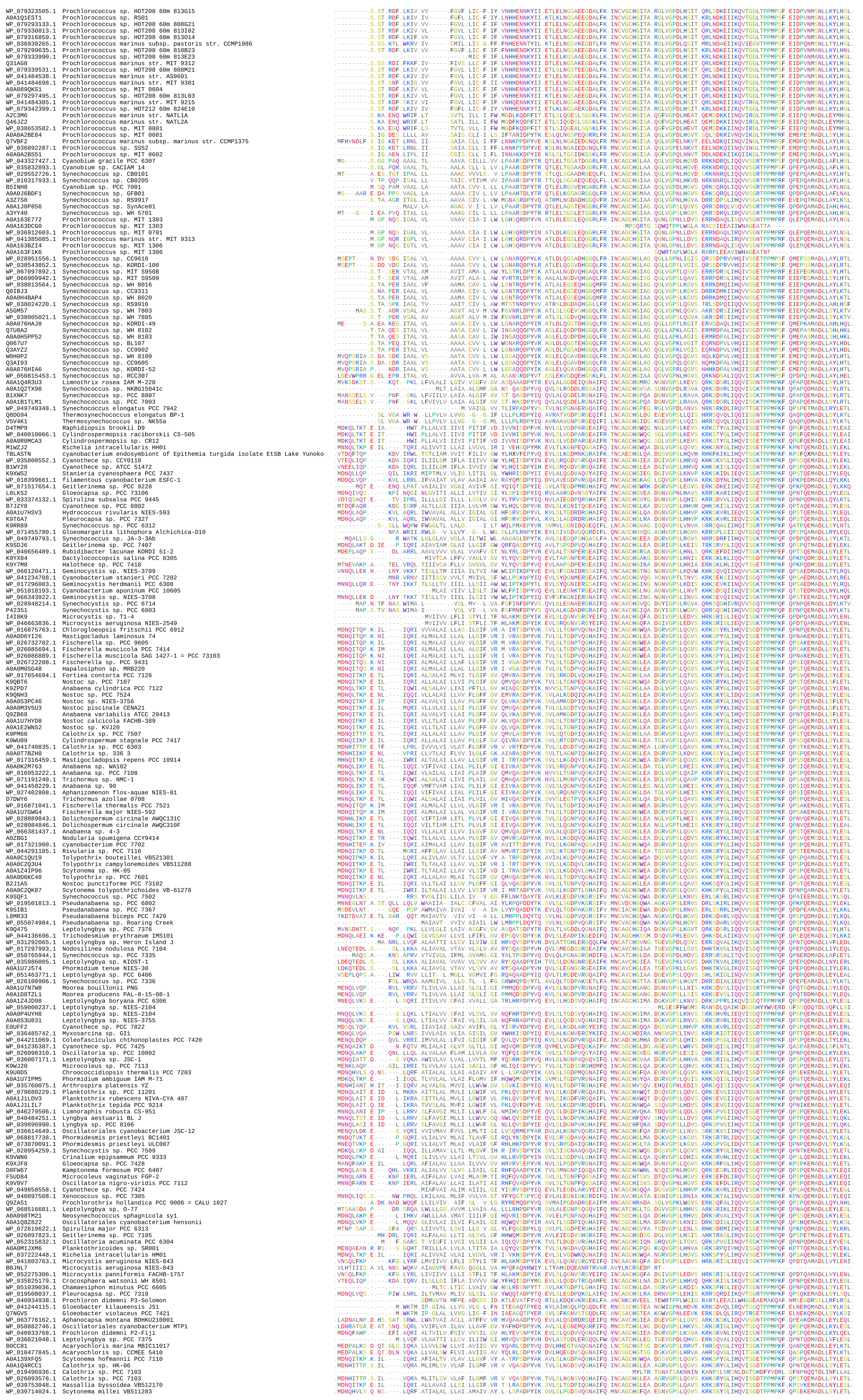
