## Supplementary material for "Cytochrome c_M_ downscales photosynthesis under photomixotrophy in *Synechocystis* sp. PCC 6803": Suppl data

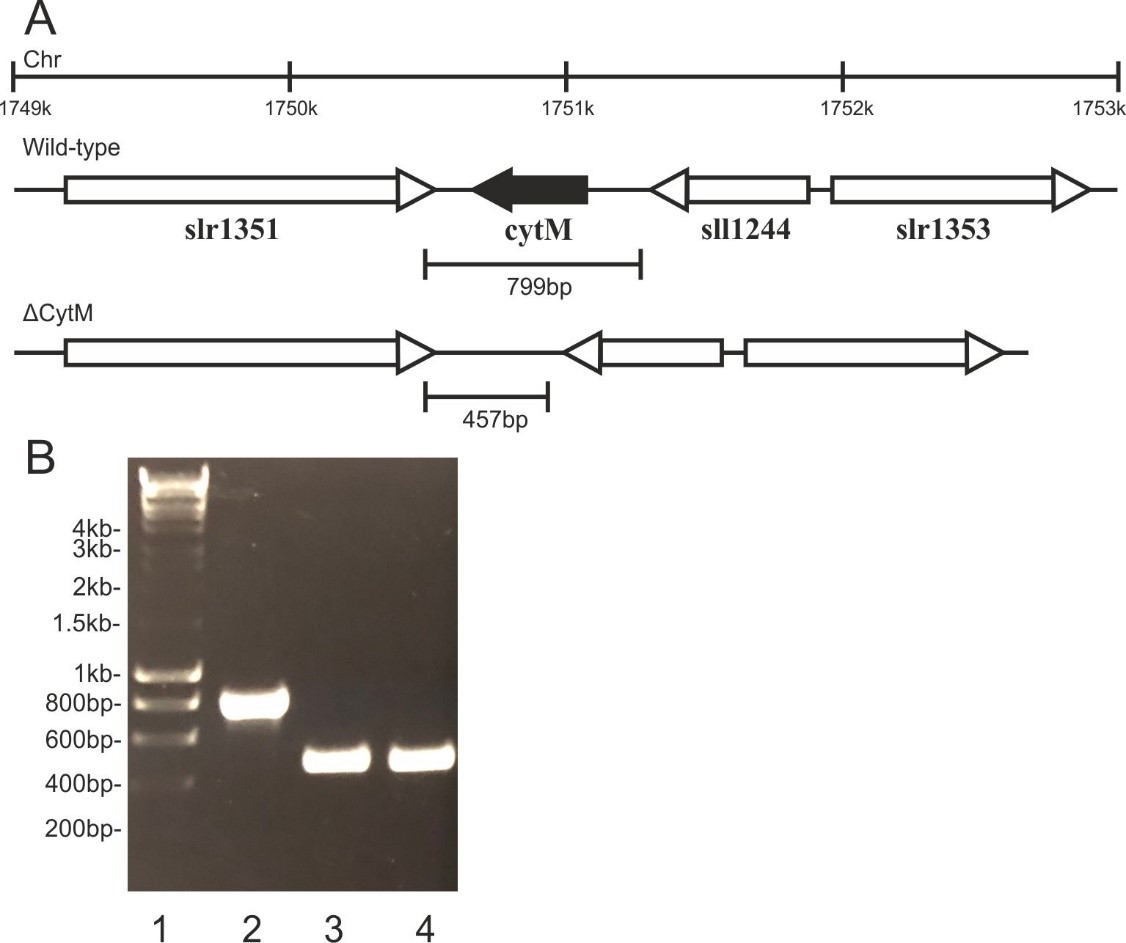
Supplemental data

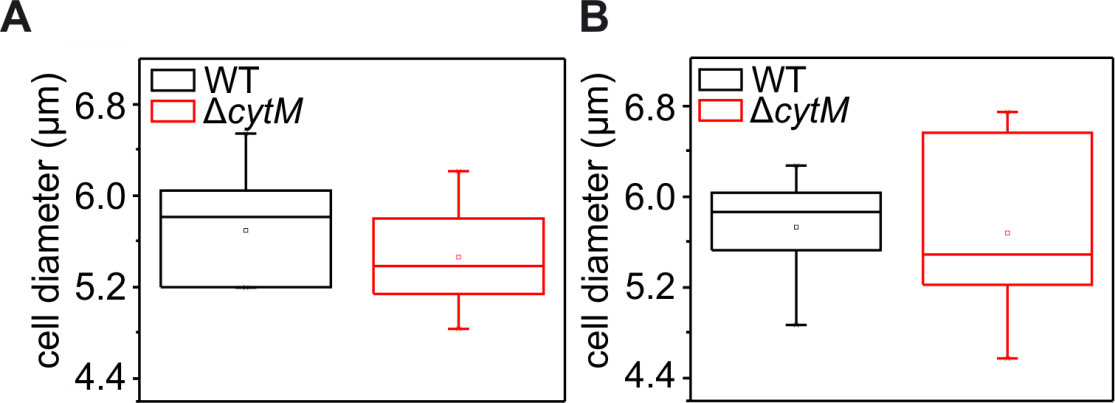
**Supplemental figure S2. Generation of *cytM* deletion mutants in *Synechocystis*.** Schematic representation of the *cytM* locus in *Synechocystis* (A) showing the genomic location (top) and profiles of the WT (middle) and the unmarked knockout mutant (bottom). Regions deleted in the mutant strain are shaded in black. Amplification of genomic DNA (B) from wild-type (lane 2), ΔCytM (lane 3) ΔCox/Cyd/CytM (lane 4) strains using CytMf and CytMr primers (Table S1). Markers are shown in lane 1.

**Supplemental figure S3.** **Cell size of WT and ΔCytM grown photomixotrophically and photoautotrophically.** Cell diameter of photomixotrophically (A) and photoautotropically (B) cultured cells was determined by microscopy and brightfield image analysis. The diameter of photoautotrophically cultured WT cells falls into the range of 1.67 - 2.46 µm (Zavřel et al., 2017). Due to the glow around the cells however, our method overestimated the cell size by a factor of 3. Nevertheless, the overestimated values are displayed rather than numbers corrected by an empirical factor. Cultivation is detailed described in Fig. 2. Values are means ± SD, n=2-3 biological repeats.

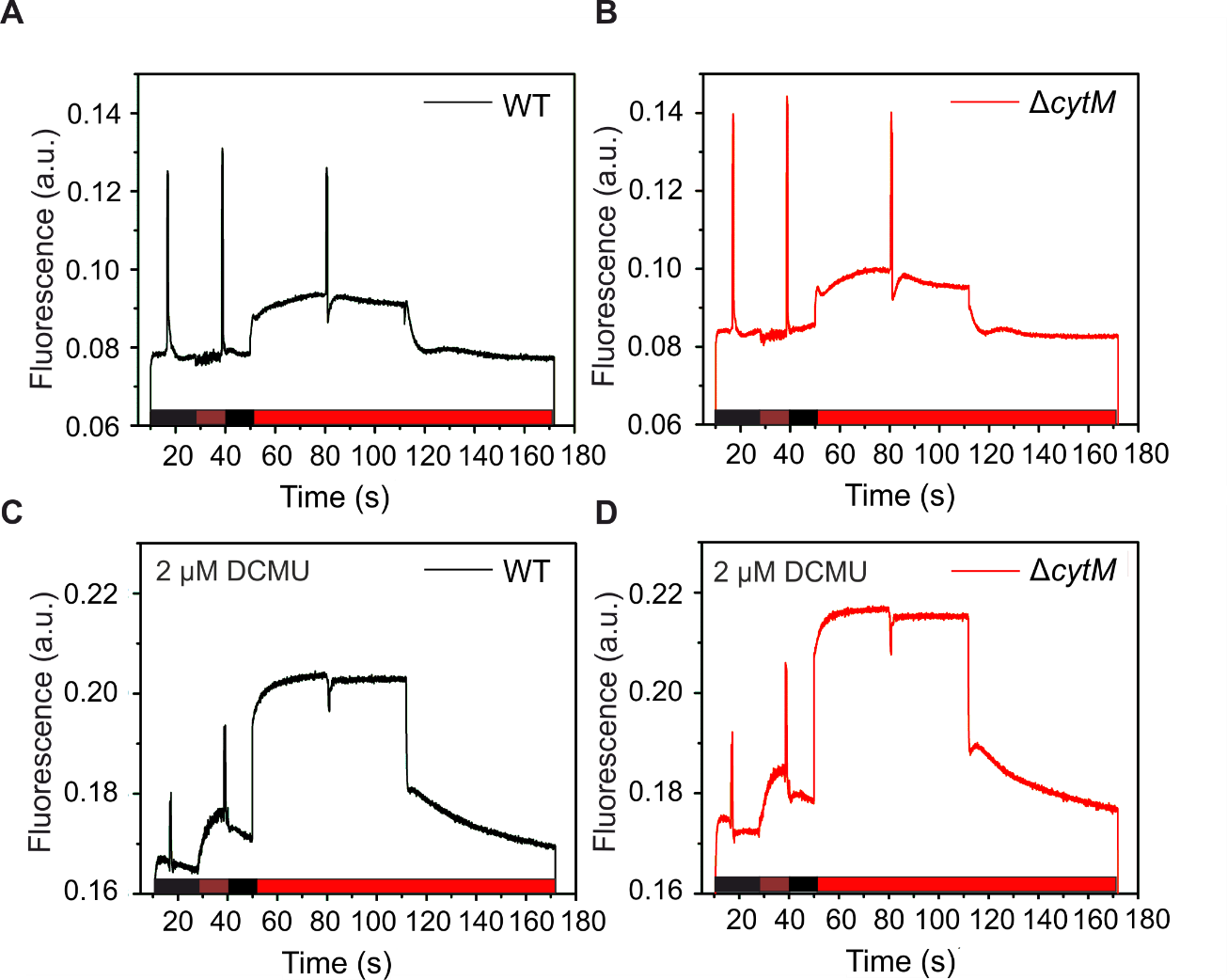
**Supplemental figure S4****. Fluorescence transients of photoautotrophically cultivated WT and ΔCytM determined in the presence of 2 µM DCMU.** Cultivation is described in Fig. 2. Prior to measurements, cell suspension was adjusted to 7.5 µg chl ml^1-^, resuspended in fresh BG-11 and dark adapted for 5 min. Fluorescence was recorded in darkness (black bars), under 40 W m^−2^ far-red light (brown bars) and under 50 µmol photons m^−2^ s^−1^ actinic red light (red bars). The graphs are representative of 3 biological replicates.

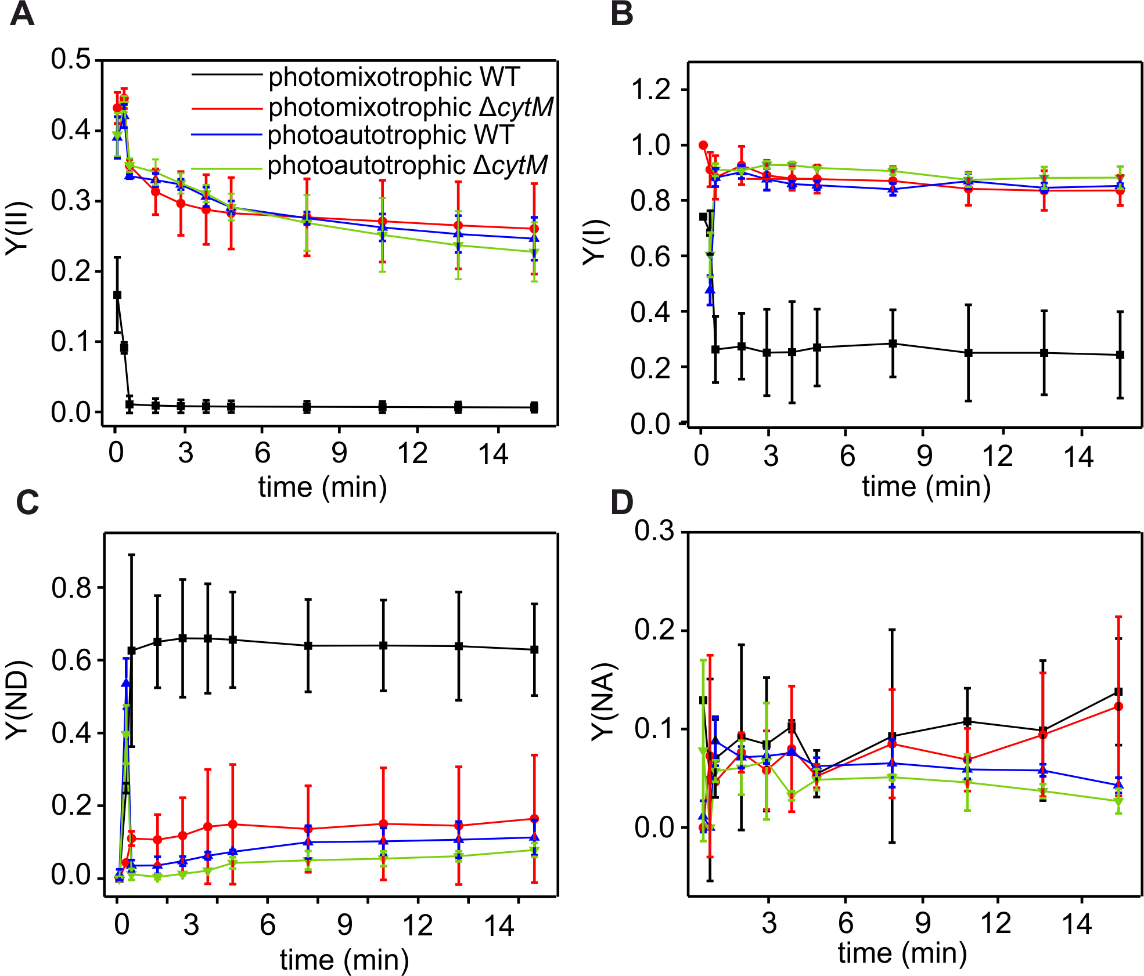

**Supplemental figure S5.** **Photosynthetic parameters of WT and ΔCytM grown photomixotrophically and photoautotrophically.** PSI yield (A), PSII yield (B), yield of PSI acceptor side limitation (C) and yield of PSI donor side limitation (D). Cultivation, sample preparation and the experimental conditions are described in Fig. 3. Parameters are calculated from the data shown in supplemental Fig. S7 and S8.

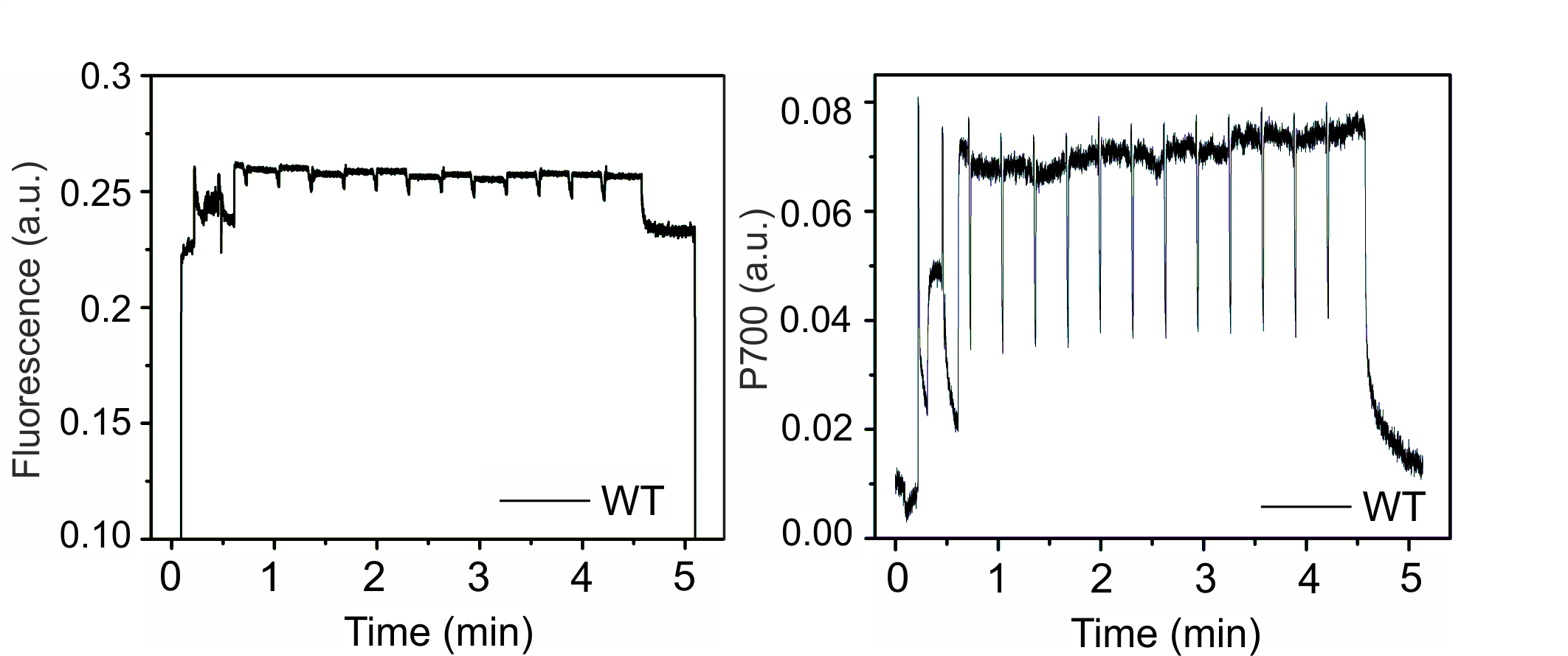

**Supplemental figure S6.** **Fluorescence transients and P700 oxidoreduction on the third day of photomixotrophic growth of WT *Synechocystis* substrain.** Cultivation, sample preparation and the experiment were carried out according to Fig. 3.

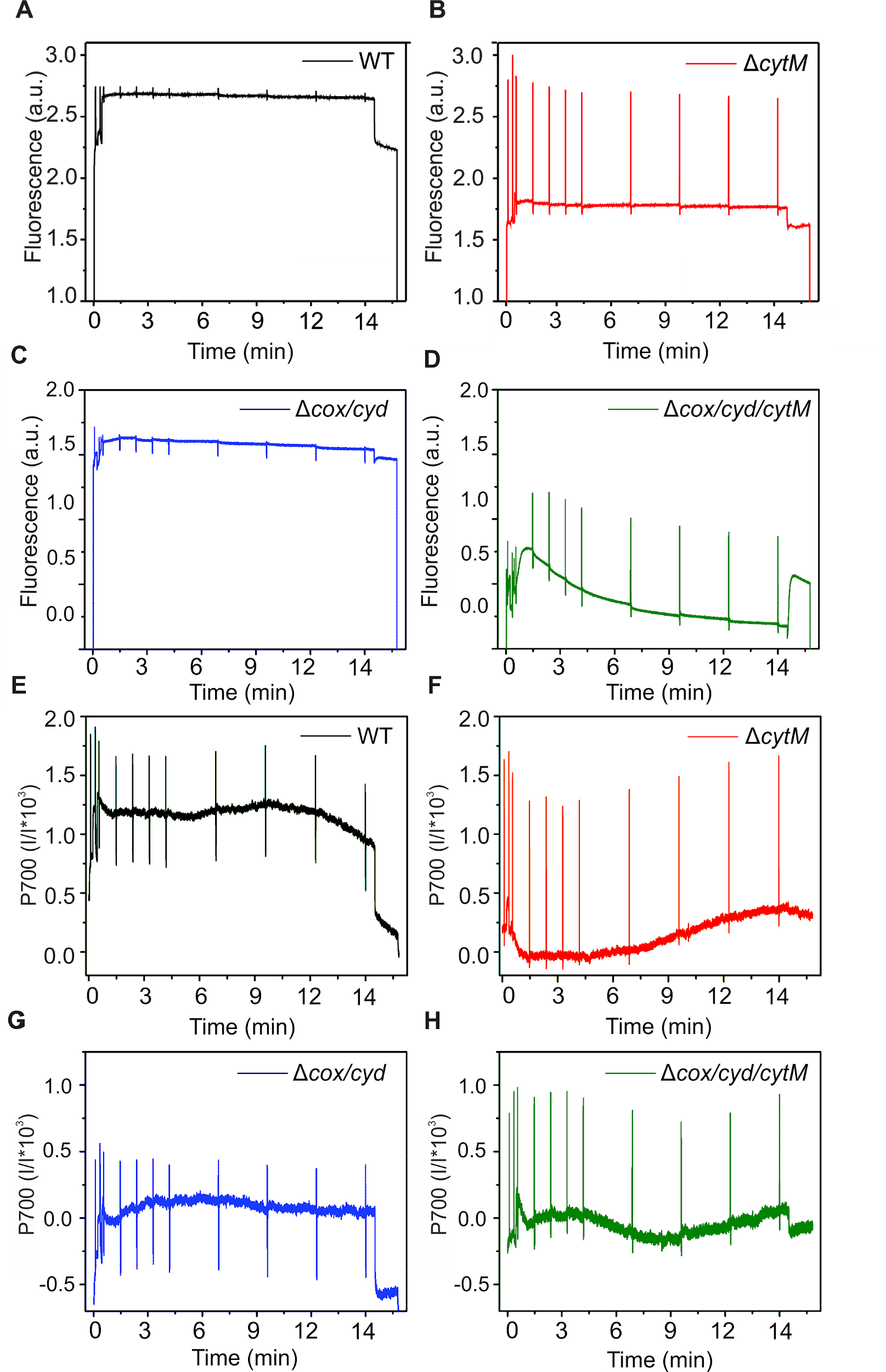

**Supplemental figure S7.** **Fluorescence transients and P700 oxidoreduction kinetics of photomixotrophically grown WT, ΔCytM, ΔCox/Cyd and ΔCox/Cyd/CytM.** Cultivation, sample preparation and the experimental conditions are described in Fig. 3.

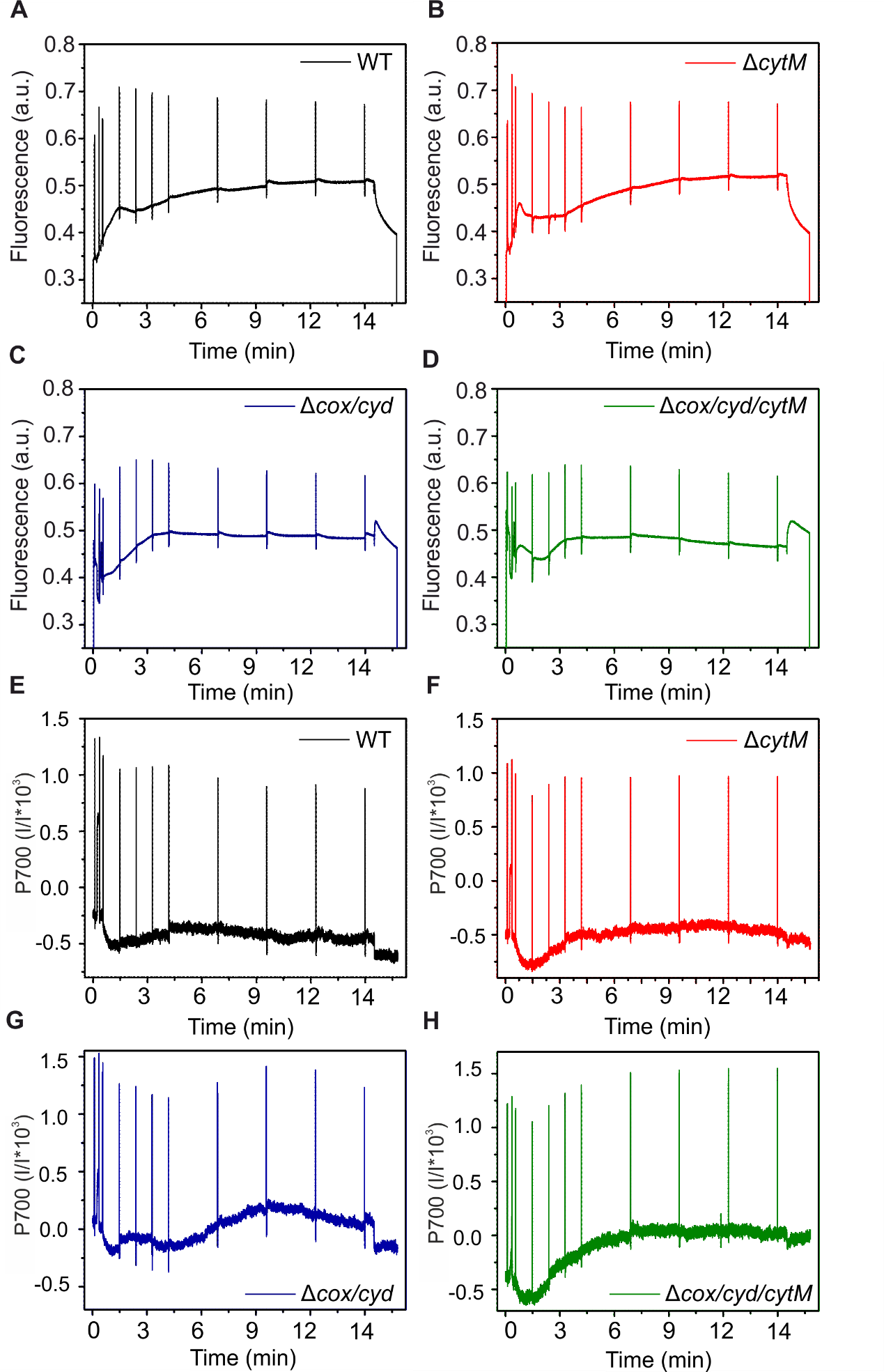

**Supplemental figure S8.** **Fluorescence transients and P700 oxido-reduxtion kinetics of photoautotrophically grown WT, ΔCytM, ΔCox/Cyd and ΔCox/Cyd/CytM.** C Cultivation, sample preparation and the experimental conditions are described in Fig. 3.

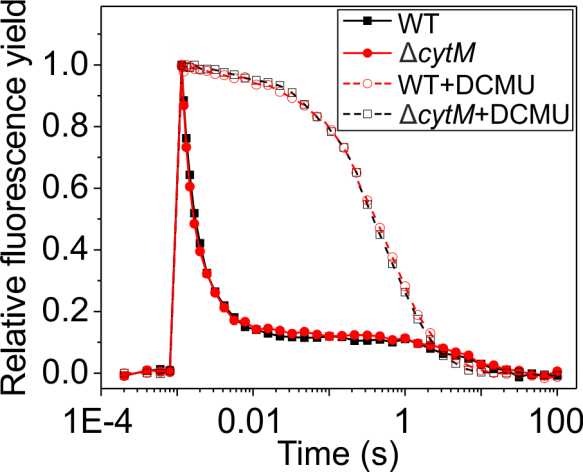

**Supplemental figure S9.** **Flash-induced increase of fluorescence yield and its relaxation in dark in photoautotrophically grown WT and ΔCytM.** Cultivation is detailed in Fig. 3. Prior to measurements, cell suspensions were adjusted to 5 µg chl ml^-1^, the samples were incubated in the dark for 5 min and DCMU was added to a final concentration of 20 µM.

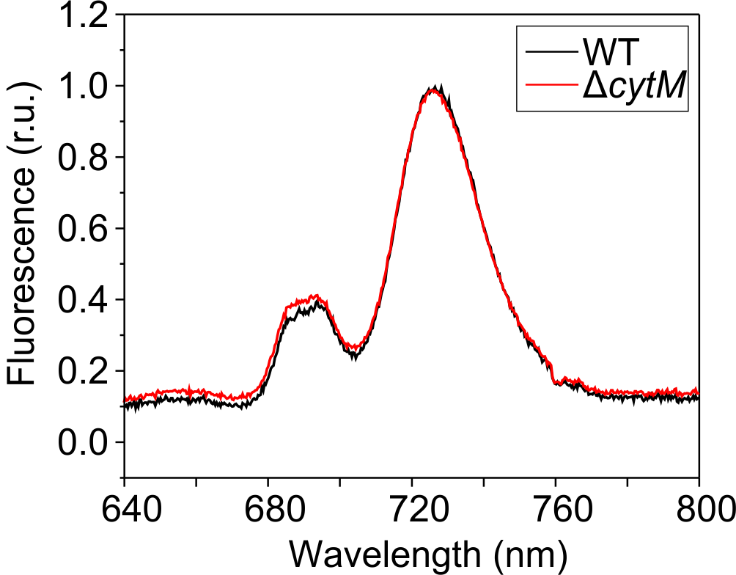

**Supplemental figure S10.** **77K steady state fluorescence emission spectra of WT and ΔCytM grown photomixotrophically.** Cultivation is detailed in Fig. 3. Prior to the measurement, samples were adjusted to 7.5 µg chl ml^−1^ and light-adapted to those conditions under which the cells were cultivated. Samples were excited at 440 nm. The curves are normalized to their respective PSI emission peak at 723 nm.

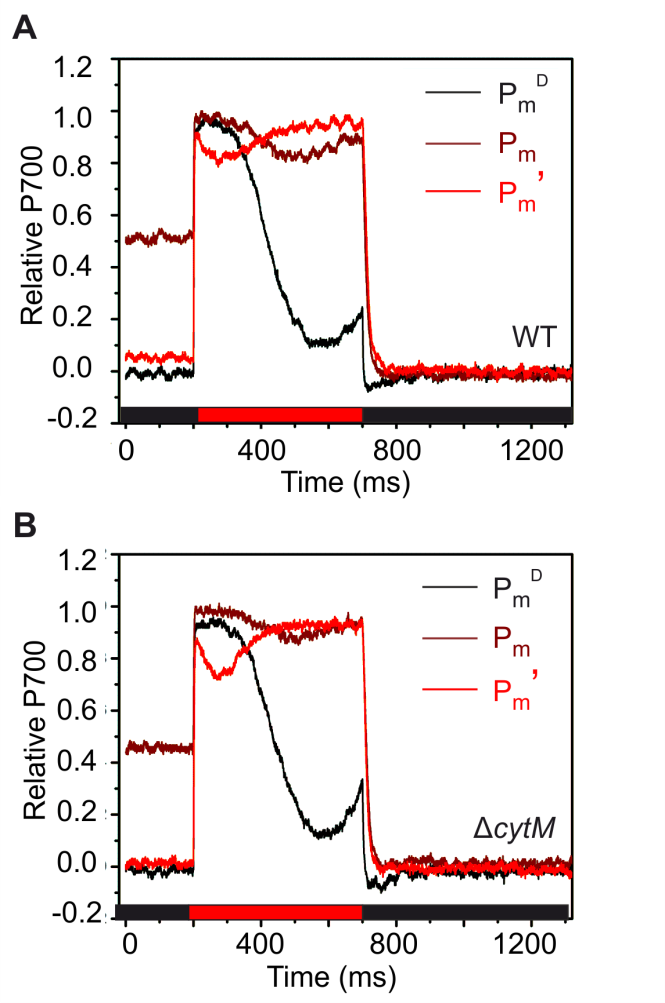

**Supplemental Fig. S11.** **Fast kinetics of** **P700 oxidoreduction of WT and ΔCytM grown under photoautotrophic conditions.** P700 fast kinetics of WT (A) and ΔCytM (B) cultivated photoautotrophically. Cultivation, sample preparation and experimental conditions are described in Fig. 3. P_m_^D^, kinetics of dark adapted cells in darkness; P_m_, kinetics under far red light; P_m_’ kinetics under actinic red light. The curves are normalized to P_0_ and P_m_.

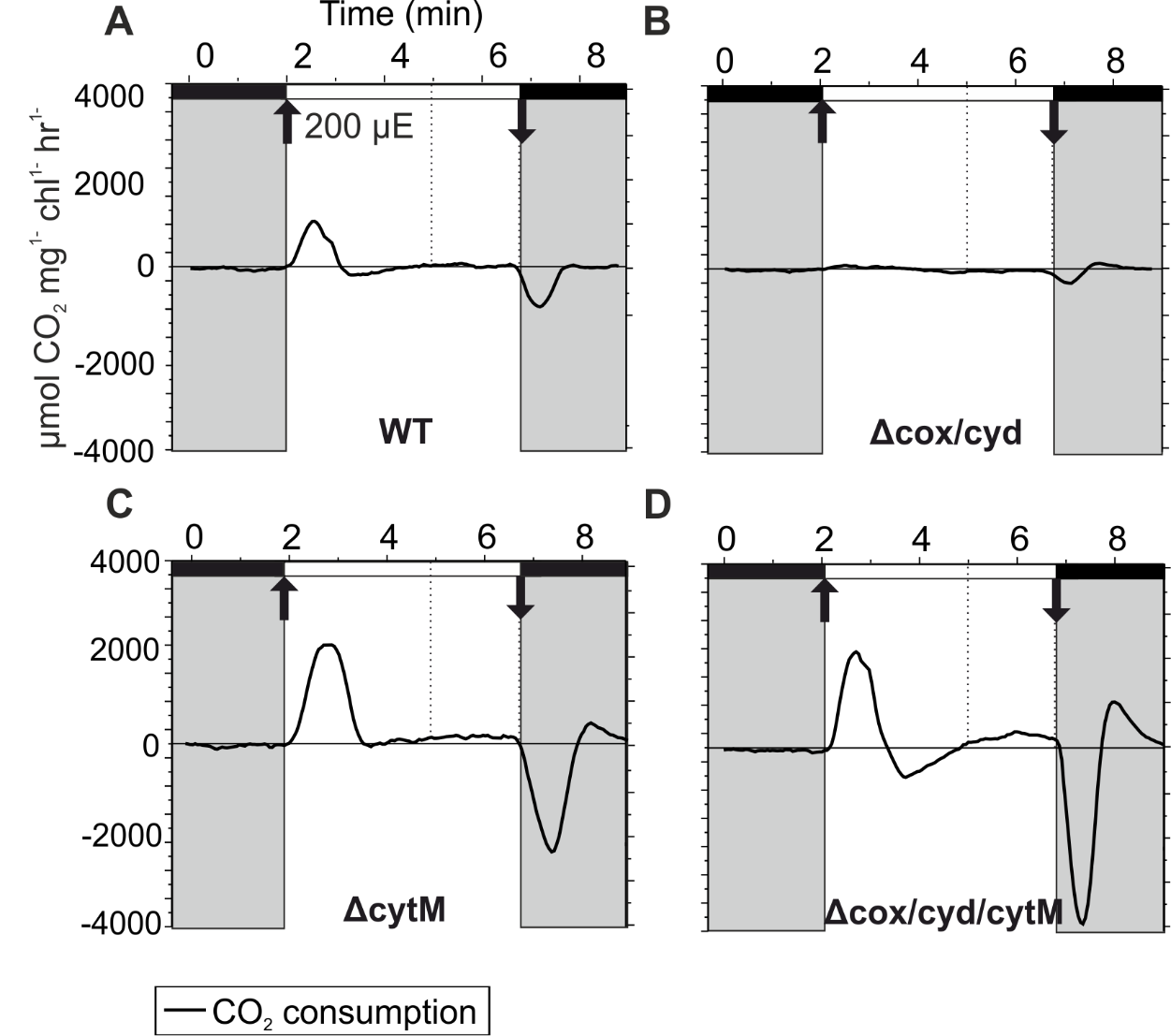

**Supplemental figure S12.** **The rate of CO_2_ flux in photomixotrophically grown WT, ΔCytM, ΔCox/Cyd and ΔCox/Cyd/CytM.** Omitted data from Fig. 6. Cultivation and sample preparation are described in Fig. 3. Dissolved C_i_ concentration was adjusted to 1.5 mM by adding NaHCO_3_. In the light phase, cells were illuminated with 200 µmol photons m^−2^ s^−1^ (200 µE) white light. The graphs are representatives of 3-6 biological replicates.

**Supplemental Table S1. List of oligonucleotides used in this study.**

| **Primer name** | **Sequence (5’ to 3’)** |
| --- | --- |
| CytMleftfor | GATCGAATTCGCCATTCTCAATCGGGATAA |
| CytMleftrev | GATCGAGCTCGGAGATGGCGGATTTATTGA |
| CytMrightfor | GATCGGATCCGGCCATGGGTTAAGAACGAT |
| CytMrightrev | GCATTCTAGAGGGATAATGTTGGCGAAAAA |
| CytMf | TTGGCAGAAAATCCCACTTC |
| CytMr | GGCCGAGCTGGATAGATTTT |

**Supplemental Table S2. Rates of O_2_ and CO_2_ fluxes in photomixotrophically grown WT, ΔCytM, ΔCox/Cyd and ΔCox/Cyd/CytM.** Source data of Fig. 6A.

|  | **WT** | **ΔCytM** | **ΔCox/Cyd** | **ΔCox/Cyd/CytM** |
| --- | --- | --- | --- | --- |
| **O_2_ consumption in dark** | 20.94±7.42 | 30.64±6.80 | 14.43±6.39 | 14.60±6.21 |
| **Gross O_2_ production** | 11.13±11.41 | 110.05±28.92 | 2.18±0.95 | 134.09±16.82 |
| **O_2_ consumption in light** | 13.38±7.28 | 47.99±13.75 | 12.18±4.57 | 38.93±12.85 |
| **Light induced O_2_ consumption** | -7.56±0.82 | 17.36±11.18 | -2.25±1.82 | 24.33±11.82 |
| **CO_2_ consumption** | -28.95±52.94 | 127.13±27.94 | -42.45±18.90 | 167.68±30.14 |
| **Net O_2_ production** | -2.26±6.53 | 62.06±32.16 | -10.00±3.88 | 95.16±19.39 |
| **Post-illumination O_2_ consumption in dark** | 18.02±6.33 | 28.93±8.44 | -8.08±28.39 | 23.47±7.84 |

Supplemental materials and methods

P_m_ determination

For maximal P700 (P_m_), the zero P700 signal level (P_0_) for fully reduced P700 is determined, so changes in the P700 signal are evaluated with respect to this reference level. The Dual PAM v1.19 software determines P_0_ as the relaxed P700 signal briefly after the saturating pulse. This is based on the assumption, that P700^+^ is fully reduced in the subsequent darkness. However, P700^+^ relaxation is markedly slower in photomixotrophically cultured WT compared to photoautotrophically cultured cells. As a result, the software overestimates P_0_ and we thus calculated P_0_ manually. P_0_ was determined as the initial P700 after 15 min dark adaptation, assuming that after long dark adaptation, the P700 pool is fully reduced. P700 signal was referenced then against P_0_ and P_m_ was determined as the maximal P700 observed.

Supplemental literature cited

Zavřel, T., Očenášová, P., Červený, J., 2017. Phenotypic characterization of Synechocystis sp. PCC 6803 substrains reveals differences in sensitivity to abiotic stress. PLoS One 12.
